# The mechanism and consequences of amyloid-β modulating thiamine pyrophosphokinase-1 expression in microglia

**DOI:** 10.1101/2024.09.18.613405

**Authors:** Xiaoqin Cheng, Ruoqi Zhao, Hongyan Qiu, Peiwen Song, Lanwen Kou, Shaoming Sang, Yingfeng Xia, Wenwen Cai, Boru Jin, Qiang Huang, Peng Yuan, Chunjiu Zhong

**Author notes:** These authors contributed equally to this work (Xiaoqin Cheng, Ruoqi Zhao, Hongyan Qiu, Peiwen Song).

## Abstract

Ample studies attribute cognitive decline in Alzheimer’s disease to amyloid-β deposition ^1–6^. However, brain amyloid-β accumulation that saturates years before the manifestation of clinical symptoms is dissociated with cognitive decline of the disease ^7^. It is unknown how these two processes are mechanistically linked. In this and our accompanied study, we report that thiamine pyrophosphokinase-1 (TPK) deficiency plays essential roles in both processes via distinct mechanisms. Here we describe that diminished microglia Tpk controls the propagation of amyloid-β plaques. In APP/PS1 transgenic mice, microglia showed elevated *Tpk* expression at 2-month-old, but reduction in a plaque-centric manner at 8-month-old. Interestingly, lipopolysaccharide, but not amyloid-β, induceed *Tpk* reduction in cultured microglia. *Tpk* reduction led to microglia dysfunction, showing volatile motility but reduced phagocytosis and weak response to focal tissue injury, with accumulation of intracellular lipid droplets and abnormal mitochrondria. In Alzheimer’s disease mice, microglia-specific knockout of *Tpk* caused diminished plaque coverage, exacerbated plaque burden and synaptic loss. However, increased plaques were not accompanied by the development of neurofibrillary tangles or brain atrophy, in contrast to the phenotype described in our accompanied paper with neuronal *Tpk* deletion. In conclusion, plaque-induced inflammation reduces *Tpk* in microglia, selectively exacerbating the spread of amyloid pathology.

---

Multiple lines of evidence support that the accumulation of amyloid-β (Aβ) plaques as an initial causal factor in Alzheimer’s disease (AD)^1,2,4–7^. Several hypotheses have been proposed regarding the direct mechanistic link between amyloid plaque and neural network dysfunction, including blockade of synaptic transmission^8,9^, interference with synapse turnover^10^, and disruption of axonal transmission^11,12^. However, these mechanisms fall short in explaining the decades of time lag between the manifestation of cognitive decline and the formation of amyloid plaques^13^. On the other hand, the accumulation of phosphorylated Tau proteins and brain atrophy show a closer correlation with cognitive decline^14–16^. Thus, distinct molecular and cellular events could constitute the processes of plaque accumulation and cognitive defect. Only a few studies have so far attempted to examine the link between these processes^17^, while the molecular mechanism remains largely unknown.

In this and our accompanied study, we show that the deficiency of a single protein, thiamine pyrophosphokinase-1 (TPK), could serve as a critical link between these two processes. TPK is an enzyme that converts thiamine into functional thiamine diphosphate (TDP), which is an essential cofactor for the respiratory chain reactions^18–21^. As described in our accompanied study, *Tpk* deficiency in the brain is a specific feature of sporadic AD, but not seen in other types of dementia. And neuronal depletion of *Tpk* prompts the development of neurofibrillary tangles and severe brain atrophy. Here, we report that Aβ plaque induces an inflammatory microenvironment that stimulates the surrounding microglia and reduces their Tpk expression. Tpk reduction leads to microglia dysfunction, eventually exacerbating the plaque pathology. Together, these studies indicate that the loss of Tpk in different types of cells promotes distinct aspects of pathological change seen in AD. Reversing the deficiency in Tpk or related proteins could therefore constitute a new class of treatment for AD.

## Result

### Plaque-associated microglia show reduced expression of TPK

In our accompanied studies, we reported that brain samples from AD patients showed significantly decreased TPK expression compared to samples from age-matched control subjects. While *TPK* plays essential roles in metabolism in many cell types, the expression level in microglia is the highest among all types of cells in the brain (http://brainrnaseq.org/)^22^. In order to understand the changes of *TPK* expression specifically in microglia associated with AD, we analyzed a previously published single-cell transcriptome dataset from human AD and control subjects^23^. We found a reduced level of *TPK* in microglia from AD patients (**Extended Fig 1**).

Having identified a reduction of *TPK* expression in the microglia of AD patients, we hypothesized that Aβ enriched in the micro-environment around plaques could be a contributing factor, and proceeded to directly test the effect of Aβ on microglia TPK expression. To this end, we isolated and purified primary microglia from neonatal mice, and treated the cells with synthetic Aβ peptides (**Fig. 1a**). Contrary to our hypothesis, we found that Aβ treatment led to an initial acute increase of TPK expression in microglia (**Fig. 1b, c**).

**Figure 1:**
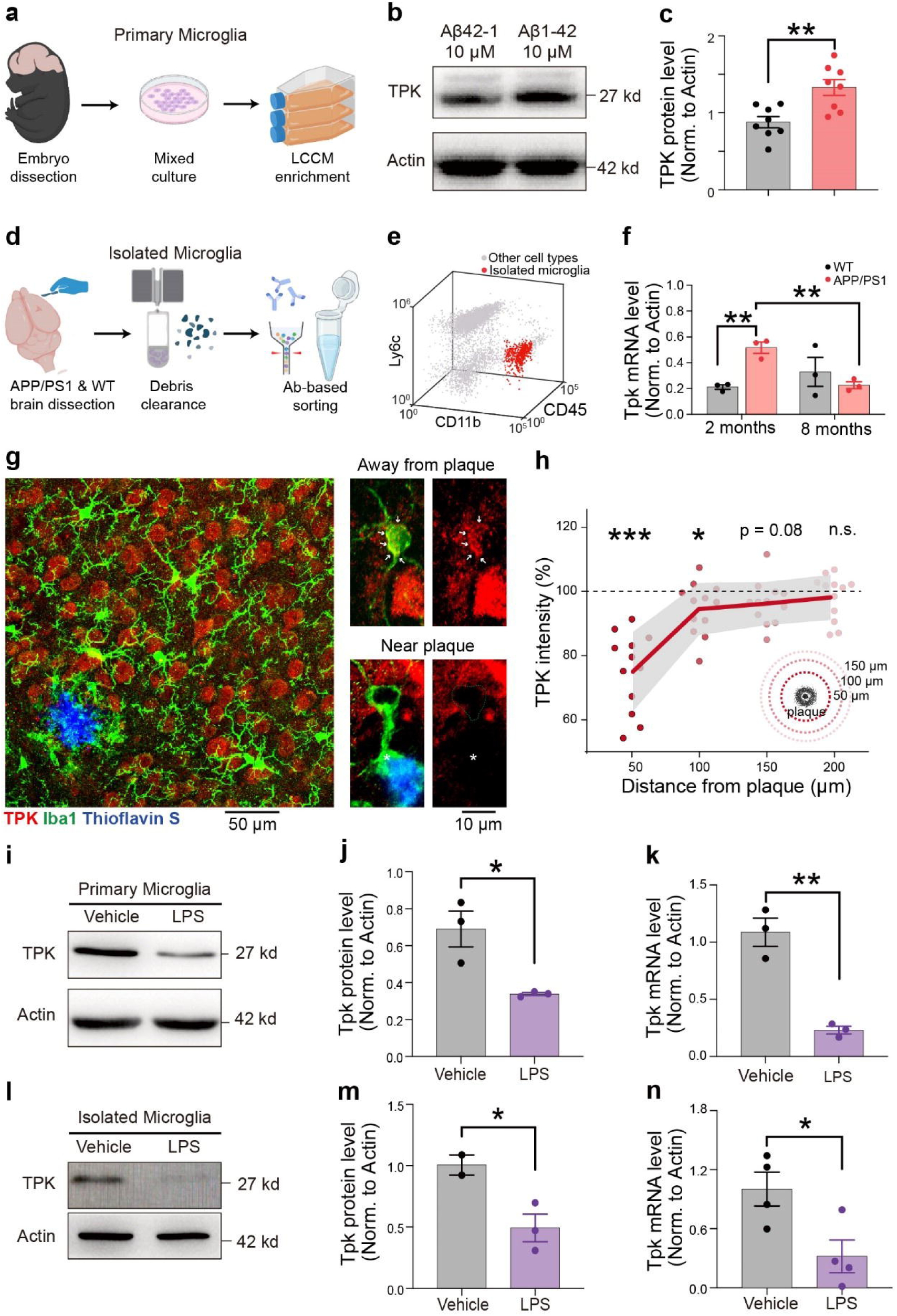
Inflammation, but not Aβ, reduces TPK expression in microglia. **a**, A diagram of primary microglia extraction from neonatal mice brain by LCCM enriched mixed glia culture. **b,c,** Western blot analysis of TPK protein levels in primary microglia treated with 10 μM of Aβ 1-42 or Aβ 42-1 for 12 hours. n = 3 independent experiments with total samples indicated in each group. Representative result image **(b)** and statistical analysis **(c)**. **d,** Flow sorting scheme for the isolation of microglia from adult APP/PS1 and wild type mice. **e,** CD11b^+^CD45^low^Ly-6c^-^ labeled cells were defined as microglia for FACS sorting. **f,** Quantifications of *Tpk* mRNA expression detected by one-step real-time quantitative RT-PCR in microglia isolated from 2 and 8-month-old APP/PS1 mice and littermate mice. nL=L3 mice per group. **g,** Representative confocal images of triple immunofluorescent staining of microglia (green), Aβ (blue) and *Tpk* (red) in cortical regions from 8-month-old APP/PS1 mice. Arrows indicate TPK expression in microglia away from plague, asterisk indicates TPK expression in microglia near plague. **h,** Quantifications of TPK intensity in microglia at different distances from plague borders in 8-month-old APP/PS1 mice, n =L12 mice per group. **i-k,** The mRNA and protein levels of *Tpk* were detected by real-time PCR (**k**) and western blotting (**i** for representative image and **j** for quantification) in primary microglia treated with 500 ng/mL LPS or equal volume of vehicle (PBS) 24 hours before sacrifice. After treatment with LPS, both mRNA and protein level of *Tpk* in primary microglia were significantly decreased, n = 3 independent experiments. **l-n,** The mRNA and protein levels of *Tpk* were detected by real-time PCR (**n**) and western blotting (**l** for representative image and **m** for quantification) in isolated microglia treated with 5 mg/kg LPS or vehicle (PBS) for 24 hours. for western blotting, n = 2 mice for vehicle group and n=3 mice for LPS group. For mRNA test, n = 4 mice per group. After treatment with LPS, both the mRNA and protein level of *Tpk* in primary microglia were significantly decreased. Statistical analyses were performed using two-tailed t-tests and one-way ANOVA in **h**. Correction for multiple comparison in **c** was done by setting a false discovery rate at 5%. Summary data represent means ± SEM. *p < 0.05, **p < 0.01, ***p < 0.001; n.s., p > 0.05.

We then investigated the change of *Tpk* expression in microglia in AD-like APP/PS1 mice. To this end, we used a flow cytometry-based cell sorting approach and selected microglia as CD11b^+^, CD45^lo^, and Ly6c^-^ cells (**Fig. 1d-e**). By quantifying the amount of *Tpk* mRNA in microglia isolated from APP/PS1 mice and compared to age-matched littermate controls, we found that microglia in APP/PS1 mice showed elevated *Tpk* expression at 2-month age, and converted to reduced *Tpk* expression at 8-month-old (**Fig. 1f**). To further examined the spatial pattern of TPK in microglia, we stained tissue sections from APP/PS1 mice with anti-TPK antibodies, and imaged the staining by high-resolution confocal microscopy. Our data showed that microglia TPK proteins primarily located in the soma regions (**Extended Fig.1**). Interestingly, we found a specific reduction of TPK staining in microglia with close proximity to amyloid plaques (**Fig. 1g-h**). This plaque-associated TPK reduction could contribute to the reduced TPK expression seen in microglia from AD patients and transgenic mice.

### Inflammation state, not amyloid, suppresses TPK in microglia

The above results indicate that TPK expression was decreased in adult APP/PS1 mice and AD patients, but Aβ peptide per se is not sufficient to induce TPK reduction in microglia. Previous research has showed that plaque-associated microglia exhibit a chronic and pathological activation state ^24^. We thus suspect that it is this activation state that causes the reduction of TPK expression. To test this hypothesis, we treated the cultured primary microglia with lipopolysaccharide (LPS) for 24 hours and examined the levels of *Tpk* mRNA and protein. We found a robust reduction of TPK expression with LPS treatment (**Fig. 1i-k**). We further examined whether the same activation could stimulate TPK reduction in microglia *in vivo*. We injected LPS intraperitoneally in mice and collected the brain after 1 day. LPS injection led to pronounced activation of microglia (**Extended Fig. 1**), and a very potent elevation in the levels of proinflammatory tumor necrosis factor-□ (TNF□) (**Extended Fig. 1**). At the same time, similar to the results from cultured microglia, we observed marked reduction of *Tpk* mRNAs as well as proteins (**Fig. 1l-n**). Together, these data indicated that the inflammation-like activation state could be contributing to the reduction of TPK in plaque-associated microglia.

### Microglia lacking *Tpk* show altered metabolism

To examine the impact of TPK reduction in microglia, we generated a conditional knockout mouse line of *Tpk* (**Extended. Fig. 1**). By crossbreeding this line with the microglia specific promoter derived Cre line (Cx3cr1-CreERT2), we could specifically delete *Tpk* in microglia (mcKO) by tamoxifen induction (**Extended. Fig. 1**). We then extracted the microglia from mcKO and control mice, and examined their transcriptome. The canonical function of *Tpk* is the key enzyme in converting thiamine into TDP, which is an important co-factor for key enzymes of glycolysis, tricarboxylic acid cycle, and pentose phosphate pathway^20^. Thus, it is likely that TPK deficiency would affect metabolism and the cell’s oxidative stress. We found numerous gene expression changes in microglia after *Tpk* deletion (**Fig. 2a**). Interestingly, the majority of the 100 genes with the largest changes showed reduced expression with *Tpk* deletion (**Fig. 2b** and **Extended Table 1**). Examining classical microglia genes with homeostatic and activated functions^25^, we found that several of the homeostatic microglia marker showed a low expression pattern, such as *Csf1*, *Tmem119*, *C1qa* and *Hexb*, without any of the activation marker showed high expression (**Fig. 2c**). This pattern of expression suggested that *Tpk* deletion did not activate microglia, and converted microglia into a “low maintenance” state that showed reduced expression of multiple homeostatic markers.

**Figure 2:**
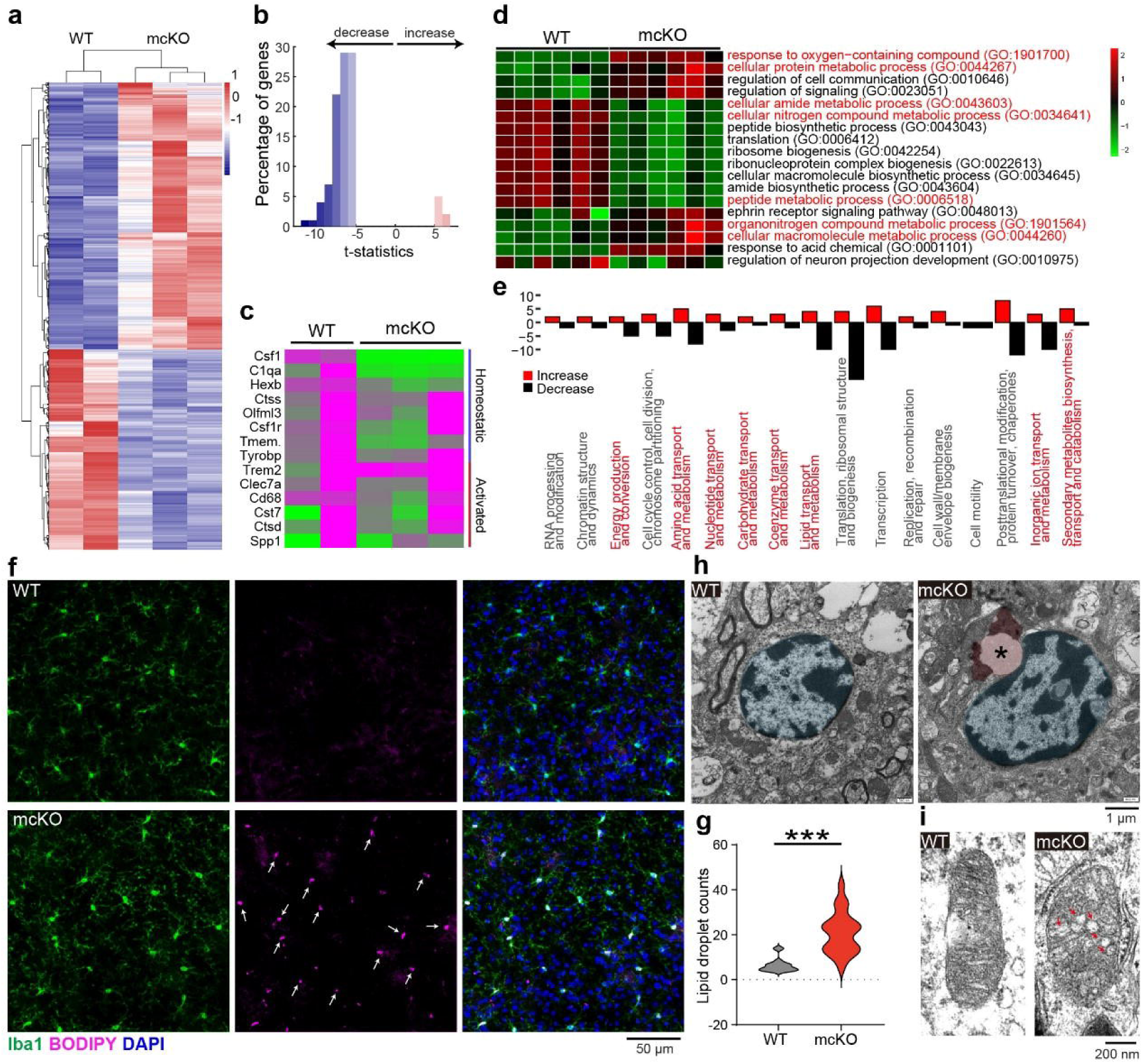
Microglia *Tpk* deletion altered their metabolic state. **a-c,** Bulk RNA-seq-based transcriptome analysis of microglia isolated from control mice or mcKO mice. Heatmaps of DEGs are shown in A. Top 100 DEGs were sub-grouped into decrease and increase group **(b)**, showing almost 90 percent of DEGs were aligned into decrease group. Heatmap showing expression level changes in homeostatic and activated microglia gene between control mice or mcKO mice **(c)**. Gene names are color coded to show their group assignments. **d,e,** Mass spectrometry analysis of brain tissues in control or mcKO mice **(d)**. GO analysis of biological functions aligning the differentially expressed genes with the EuKaryotic Orthologous Groups labels showed bidirectional modulation of expressions **(e)**. Functional groups related to oxidative stress and metabolism labeled in red. n = 6 mice per group. **f,g,** Representative confocal images of BODIPY labeled lipid droplets and Iba-1 labeled microglia in the cortical of 6-month-old mcKO mice and littermate mice, *n*L=L4 mice per group. Arrows indicate BODIPY labeled lipid droplets**(f)**. Quantification of percentage of BODIPY positiveLmicroglia in the cortical of 6-month-old mcKO mice and littermate mice**(g)**. Two-tailed t-test, ***: p<0.001. **h,** Representative electron microscope images of microglia from 6-month-old mcKO mice and littermate mice. Cell nuclear was shaded in blue and lipid droplet in red. Asterisk indicates lipid droplet. **i**, Representative electron microscope images of microglia mitochondria from 6-month-old mcKO mice and littermate mice. Arrows indicate abnormal vacuole-like mitochondria ridges.

We further analyzed the pattern of expressed proteins in mcKO mice using mass spectrometry. Consistently, GO analysis revealed alterations in the expression of multiple pathways related to oxidative stress and metabolism (**Fig. 2d**). Notably, *Tpk* deletion showed elevated expression in “response to oxygen-containing compound”, and reduced levels in multiple pathways related to metabolism. Several increased metabolism pathways suggest that perhaps *Tpk* deletion shifts microglia into a compensatory metabolic state. Consistent with this notion, aligning the differentially expressed genes with the EuKaryotic Orthologous Groups labels showed bidirectional modulation of multiple metabolism-related gene groups (**Fig. 2e**).

In addition to these changes with transcriptome, we also noticed that microglia with *Tpk* deletion showed enriched deposition of lipid droplets inside the cells, revealed by staining with the lipophilic BODIPY dye (**Fig. 2f-g**). This observation was further confirmed with electron microscopy prepared from mcKO mice brain tissue, with lipid droplets showed characteristic vacuoles-like morphology with some periphery electron dense staining (**Fig. 2g**), consistent with previous reports^26^. The elevated presence of lipid droplets has been previously associated with stressed condition in microglia ^26^. Furthermore, we frequently observed abnormal mitochondria morphology in mcKO microglia, showing dis-organized internal ridges (**Fig. 2i**). Together with the changes in transcriptome we observed, these data demonstrated that microglia metabolism was affected by *Tpk* deletion, leading to an altered metabolic state that featured lipid accumulation and diminished activation.

### *Tpk* reduction leads to volatile microglia process dynamism

We next examined the functional impact of microglia *Tpk* deletion. To this end, we crossbred the mcKO mice with a cre-dependent reporter mouse line Ai9^27^, and performed *in vivo* imaging of the labeled microglia in the mouse’s cortex. Microglia processes showed stereotypical dynamism with constant protrusions and retractions, while the soma region remained relatively stable. These dynamic movements are considered to be the basis for microglia to constantly surveil surrounding brain environment^28^. By contrasting the labeled structure at consecutive timepoints, we were able to quantify this dynamism (**Fig. 3a-b**). We found that microglia lacking *Tpk* showed elevated levels of extending and retracting processes (**Fig. 3e-f**). Consequently, we observed that *Tpk*-deleted microglia exhibited cell size enlargement and markedly elevated number of branches (**Fig. 3c-d**). These changes could be a compensatory mechanism for microglia to collect more nutrients from the environment due to *Tpk*-deletion-induced metabolic changes.

**Figure 3.**
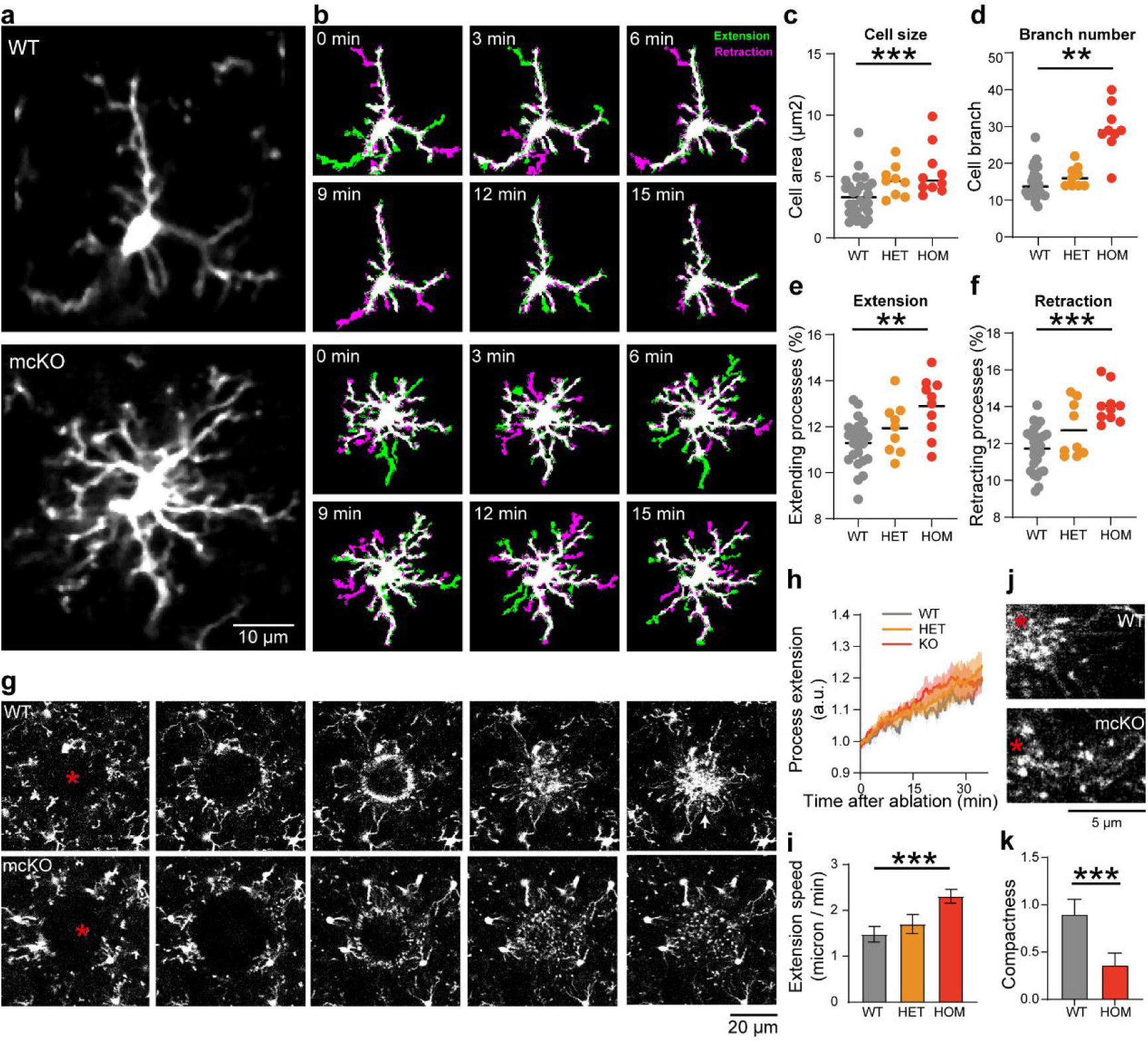
*Tpk* deficiency interferes microglia morphology and process motility. **a, b,** Representative images from in vivo two-photon timelapse imaging of microglia (Ai9 signal, white) in control (upper panel in **a** & **b**) and mcKO (lower panel in **a** & **b**) mice brains with process extension (green) and retraction (purple) at 0, 3, 6, 9, 12 and 15 minutes. **c-f,** Quantification of cell size (**c**), branch number (**d**), process extension (**e**) and retraction (**f**) of microglia in control, heterozygote and mcKO mice brains (nL=L30, 9 and 10 cells in each group, respectively). **g**, Representative imges from in vivo two-photon imaging of microglia dynamic response (Ai9 signal, white) in control (upper panel) and mcKO (lower panel) mice brains from 0 to 30 minutes after the induction of local laser ablation. Asterisks indicate the site of local laser ablation. **h**, Quantification of kinetics of microglial extension processes over 30min among different groups. The ability of extension processes is stronger in mcKO group than in control group. (nL=L4 mice) **i,** Quantification of extension speed of microglia among control, heterozygote and mcKO mice. (nL=L4 mice) **j,** Representative image of wrapping capability of microglia in contol mice (upper panel) and mcKO mice (lower panel) 30 minutes after laser ablation. Asterisks indicate the site of local laser ablation. Microglia in control group formed a tight wrapping around the injured site, whereas microglia in mcKO mice did not form tight wrapping around the injured site. **k,** Quantification of microglia wrapping compactness among control, heterozygote and mcKO mice. (nL=L4 mice) Statistical analyses were performed using one-way ANOVA. Summary data represent means ± SEM. * p < 0.05; ** p < 0.01; *** p < 0.001; n.s., p > 0.05.

An important function of microglia in the brain is the capability of responding to acute stimulants, including infection, cell debris or tissue damage^29,30^. In order to examine whether *Tpk* deletion changes the reactive functions in microglia, we performed *in vivo* laser ablation of brain tissue and monitored chemotaxis attraction of microglia to the injured site. As reported in previous studies, normal microglia showed a stereotypical directed process extension to the injured site ^28,31^ (**Fig. 3g**). Compared to normal microglia, *Tpk* deleted microglia showed increased speed of this directed movements (**Fig. 3h** and **i**), consistent with increased dynamism observed in the resting state. Interestingly, we found that *Tpk* deleted microglia did not form tight wrapping around the injured site, despite that the processes reached the site rapidly after ablation (**Fig. 3j** and **k**). These findings indicated that while the molecular machinery related to microglia process dynamism remained intact, *Tpk* deletion disrupted the capability of microglia processes to form robust and specialized wrapping, suggesting that the microglia response to pathological stimulants could be ineffective.

### *Tpk* deficiency causes ineffective plaque coverage by microglia and accelerates plaque depositions

We previously reported a protective function of microglia around amyloid plaques that forms a barrier and insulates surrounding neural tissue from the plaque’s toxicity^11,32,33^. The above results suggested that *Tpk* deletion in microglia could lead to less robust response to pathologies in the brain, we thus speculate that reduced expression of TPK in plaque-associated microglia could exert a detrimental effect and reduce the microglia coverage. We then examined the interaction between TPK expression and amyloid pathology in the brain. To this end, we first examined whether *Tpk* deletion in microglia altered the production and clearance of Aβ in the brain. Western blots and ELISA from brain extracts showed no significant changes in the mouse endogenous Aβ contents or the β-secretase necessary for its production (**Extended Fig. 2**). In addition, when we measured the phagocytic activities from cultured primary microglia, we found that *Tpk* deletion led to suppressed phagocytic activity in microglia (**Extended Fig. 2**), although at baseline state this change did not significantly alter the levels of Aβ in the brain.

In order to directly examined the impact of TPK on the interaction between microglia and amyloid, we crossbred the mcKO mice with the APP/PS1 transgenic mice to obtain APP/PS1/mcKO mice. The mice allowed us to image the plaque-microglia interaction with high-resolution microscopy. Consistent with our hypothesis, we found a reduced reactivity in plaque-associated microglia with *Tpk* deletion (**Fig. 4a-c**). *Tpk* deletion caused diminished recruitment of microglia around plaques, and the protective microglia coverage was also reduced. Our previous work proposed that this protective wrapping of plaque limits the spread of plaque pathology and its associated toxicity ^34,35^. We then measured the plaque deposition and synapse loss in APP/PS1/mcKO mice. Compared to age-sex-matched control APP/PS1 mice, we found that *Tpk* deletion in microglia caused a marked elevation in the amount of plaque deposition (**Fig. 4d-f**), which is accompanied by the exacerbated synapse loss (**Fig. 4g-h**). At the same time, these changes were not accompanied by increase in phosphorylated Tau, or neuronal loss (**Fig. 4i-o** and **Extended Fig. 3**). Taken together, these data indicate that TPK in microglia is a potent modulator that limits the expansion and toxicity of amyloid pathology.

**Figure 4:**
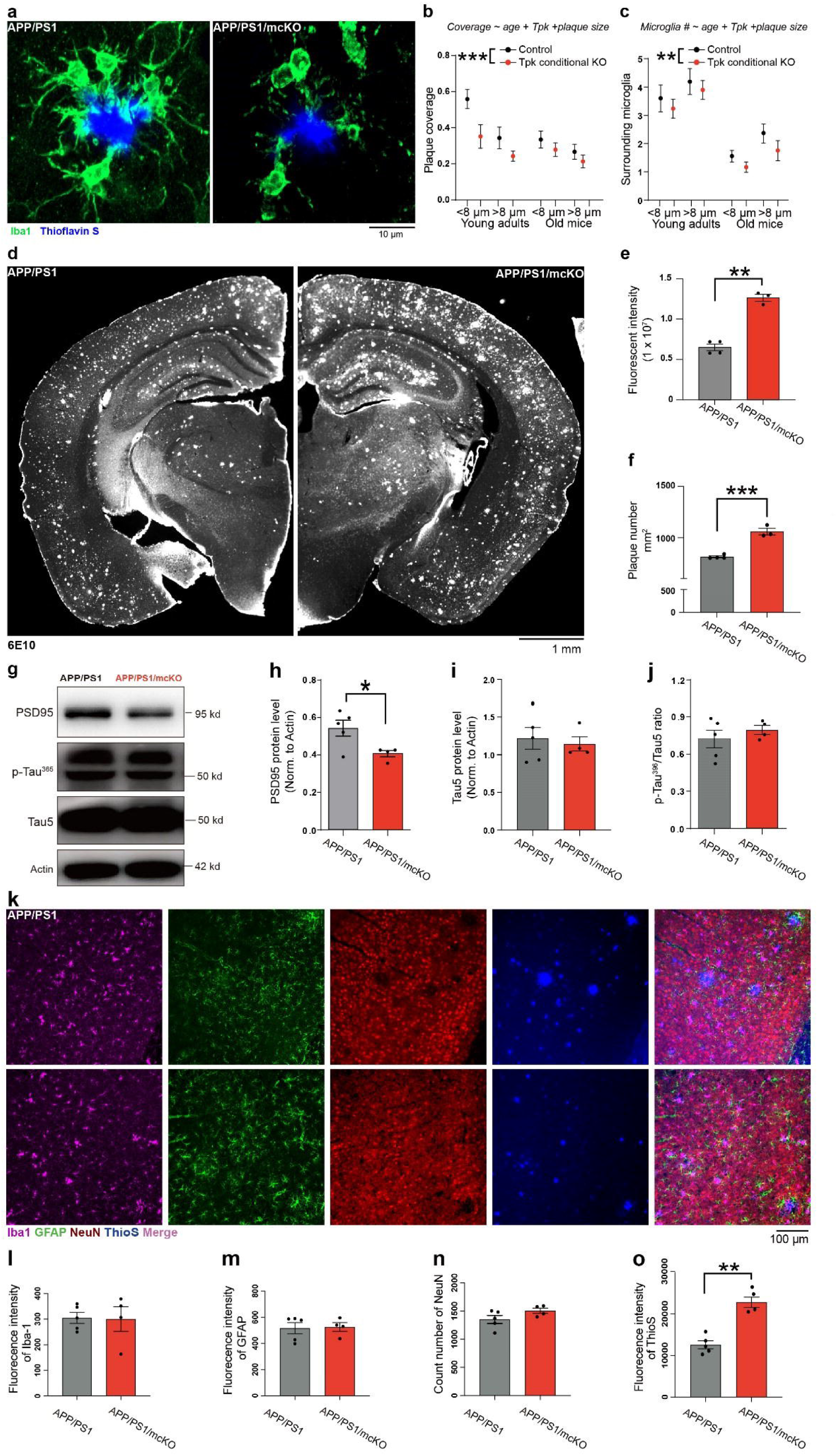
Microglial *Tpk* deletion accelerates plaque deposition but does not affect neurodegeneration. **a,** Representative confocal images of Iba-1 immunolabeled microglia coverage around Thioflavin S labeled amyloid plaques in cortical samples of 18-month-old mcKO-APP/PS1 mice and littermates. **b-c,** Quantification of the percent of plague coverage **(b)** and surrounding microglia number **(c)** among different age-matched APP/PS1 mice and mcKO-APP/PS1 mice. Eight-month-old mice were referred to young adults and 18-months referred to old mice. Mice number: 8-month-old APP/PS1 mice: n=3; 8-month-old mcKO-APP/PS1 mice: n=5; 18-month-old APP/PS1 mice: n=4, 18-month-old mcKO-APP/PS1 mice: n=3. **d-f,** Representative fluorescent staining image (**d**) and quantification of fluorescen indensity (**e)** and plaque number per 1 mm^2^ (**f**) of 6E10 immunolabeled plague in cortical samples of mcKO-APP/PS1 and their littermates. Plaque coverage was significantly heavier in APP/PS1 mice with *Tpk* conditional knockout than that in control. **g-j,** Representative western blotting images (**g**) and quantification of PSD95 **(h),** Tau5 **(i),** and p-Tau^396^/Tau5 **(j)** protein levels in brain tissue of 18-month-old mcKO mice and their littermates (nL=L4 : 5 mice). **k-o**, Representative confocal images (**k**) and quantifications of microglia number (**l**), astrocyte number (**m**), neuron number (**n**) and amyloid plaques (**o**) in brain tissue of 18-month-old mcKO mice and their littermates (nL=L4 : 5 mice). Statistical analyses were performed using unpaired two-tailed t-tests and generalized linear model (**b and c**). Summary data represent means ± SEM. Animal number per group is listed in statistical graphs. * p < 0.05, ** p < 0.01, *** p < 0.001.

## Discussion

Our current study demonstrated a vicious cycle between the formation of amyloid plaque and the deficiency of Tpk expression in microglia. The inflammatory microenvironment around amyloid plaques suppresses the Tpk expression in plaque-associated microglia. Tpk depletion is expected to interfere with intracellular glucose metabolism, blocking reactions for citric acid cycle and lipid gluconeogenesis. Unlike the rapid cell death seen in neurons (see accompanied study), microglia lacking Tpk can still survive on active glycolysis, although with limited energy resources. This alteration in microglia metabolism suppresses microglia’s function in limiting plaque growth (**Fig. 4**), as expected from previous studies ^33,36,37^. Therefore, reduced Tpk expression in plaque-associated microglia further promotes the expanding and spread of amyloid plaques.

Interestingly, the Tpk reduction in microglia was not induced by Aβ *per se*, but by the inflammatory microenvironment. Surprisingly, soluble Aβ even showed stimulating effect for Tpk expression, consistent with elevated Tpk levels in APP/PS1 mice before the onset of amyloid plaques. This raises an interesting possibility that the detrimental effect of amyloid plaques can be at least partially due to the reduction of soluble Aβ in the interstitial fluids. Several lines of evidence also support this view. For example, a longitudinal clinical study among familial AD patients showed that higher soluble Aβ42 level in CSF is negatively correlated with disease progression ^38^. Südhof, et al. further demonstrate the physiological effects of soluble Aβ in promoting synaptic genesis and transmission ^39^. In addition, drugs that lead to the reduction of soluble Aβ accelerate, rather than ameliorate, cognitive decline in individuals with AD ^40^. Recently developed Lecanemab is an antibody targeting Aβ protofibrils and delays cognitive decline of early AD patients at the endpoint of 18 months with limited but significant difference as compared with placebo ^41^. Unlike other anti-Aβ antibodies, Lecanemab significantly increases the level of soluble Aβ42 in CSF while reducing brain Aβ plaques. The effect of restoring levels of soluble Aβ on various physiological functions warrants future studies.

Importantly, *Tpk* defect in microglia in wildtype or APP/PS1 background does not lead to abnormal Aβ production, Tau hyperphosphorylation, or losses of neurons in the brain (**Fig. 4** and **Extended Fig. 2**). This is in contrast with the results seen in neuronal *Tpk* deletion, as described in our accompanied study. These results indicated clear dissociable molecular pathways for plaque accumulation and neurodegeneration. The fact that the single factor Tpk being a key control point in both these two processes places it at a unique position in AD pathogenesis. Therefore, a general deficiency of Tpk could in theory lead to all aspects of AD pathologies; and reversing this deficiency may provide therapeutic benefit. What could be the factor causing the general deficiency of Tpk? While the specific factor is currently unknown, it is likely related to glucose and lipid metabolism due to the canonical function of Tpk. Tpk deficiency may be an adaptive response to cumulative metabolic stress. Interestingly, factors related to metabolic changes often show correlation with the risk of developing AD, such as diabetes ^42^ and apolipoprotein (ApoE) ε4 isoforms ^43^. Understanding the cell type-specific mechanism of Tpk regulation could shed light on the process of AD pathogenesis.

In summary, our studies show that soluble Aβ and Aβ plaques play the opposite roles in modulating TPK expression in microglia, in which soluble Aβ enhances TPK expression in microglia, while Aβ plaques reduce its expression by inflammation. *Tpk* deletion in microglia impairs the phagocytosis and targeted process activation of microglia and thus aggravates Aβ pathology, but does not induce neurofibrillary tangles and neuron loss. Together with our accompany study, Tpk deficiency could be the underlying cause for the complete set of pathological changes seen in AD. Our studies offer mechanistic implications on the therapeutic benefits of Tpk modulation for treating AD.

## Acknowledgements

The results published here are in part based on data obtained from the AD Knowledge Portal (https://adknowledgeportal.org). Samples for this study were provided by the Rush Alzheimer’s Disease Center, Rush University Medical Center, Chicago. Data collection was supported through funding by NIA grants P30AG10161, R01AG15819, R01AG17917, R01AG30146, R01AG36836, U01AG32984, U01AG46152, the Illinois Department of Public Health, and the Translational Genomics Research Institute." Cite: https://www.nature.com/articles/s41586-019-1195-2.

We thank Dr. Yuqiu Zhang for providing the *Cx3cr1^CreERT2/+^* transgenic mice. We thank Dr. Min Jiang and Molecular and Cellular Imaging Facility of Institutes of Brain Science (IOBS), Fudan University, for their support in confocal microscopy operation, and Dr. Xueling Liao and Instumental Analysis Center school of pharmacy, Fudan University, for their support in microglial FACS sorting. This study was supported by grants from the Shanghai Municipal Science and Technology Major Project, the National Natural Science Foundation of China grants (81870822, 32371036, 91332201, 81901081, 81600930), the Shanghai Pilot Program for Basic Research – Fudan University 211TQ1400100 (22TQ019), Shanghai Natural Science Foundation (22ZR1415000), and the Natural Science Foundation of Fujian Province (2020CXB049).

## Author contributions

Xiaoqin Cheng and Peng Yuan were responsible for the experimental design and data analysis. Xiaoqin Cheng, Ruoqi Zhao and Hongyan Qiu were responsible for animal and cell experiments, mass cytometry, RNA-seq and electron microscopic experiments. Peiwen Song was responsible for in vivo imaging with two-photon microscopy and immunofluorescent staining. Lanwen Kou was responsible for re-analysis of published single-nucleus RNA-Sequencing and SMART-seq. Yingfeng Xia, Wenwen Cai and Boru Jin were responsible for Western blot and other molecular biology experiments. Shaoming Sang assisted verification and statistical analyses of the results. Peng Yuan and Chunjiu Zhong designed the study and wrote the manuscript.

## Competing interests

Chunjiu Zhong, one of the corresponding authors, holds shares of Shanghai Raising Pharmaceutical Co., Ltd., which is dedicated to developing drugs for the prevention and treatment of AD. The other authors declare that they have no competing interests.

