## Supplementary material for "The mechanism and consequences of amyloid-β modulating thiamine pyrophosphokinase-1 expression in microglia": Methods

**Materials and Methods**

1. **Analysis of human microglia TPK expression with public datasets**

For analyses the microglial TPK expression in human brain, we apply for published single-nucleus RNA-Sequencing data from The Single-cell transcriptomic analysis of Alzheimer’s disease (snRNAseqPFC_BA10) Study in Religious Order Study (ROS) cohort Religious Order Study (ROSMAP), which is available at Synapse (<https://www.synapse.org/#!Synapse:syn18485175>). The dataset contained a total of 48 human brain specimens, wherein 24 individuals diagnosed with AD as elevated β-amyloid and other pathological hallmarks, and 24 age-matched controls with no or very low Aβ burden or other pathologies.

R's Seurat program was employed. Nuclei with a mitochondrial content of more than 5% were eliminated for quality control. Two procedures were used to screen out duplets and multiplet nuclei. The first step was to eliminate any nuclei that expressed multiple marker genes. After that, cells with high UMI and gene counts per cell were eliminated.

Prior to clustering, principal component analysis was carried out, and the ElbowPlot was used to choose the first ten principal components. The FindClusters function, which operates on the K-nearest neighbor (KNN) network model with the granularity ranging from 0.4 - 1.2 and picked 0.5 for the downstream clustering, was used to accomplish the clustering. We used differential expression to find both negative and positive markers for each cluster in order to discover the markers for each cluster relative to all other clusters. To further examine the sub-clusters within each cell type, nuclei from broad cell types (astrocytes, microglia, and oligodendrocyte clusters) were extracted and re-clustered.

On the UMAP map, we located the class group of the gene we were interested in *TPK*, and using the violin map, we characterized the location of TPK1 in microglia.

2. **Animals**

All animal care and experimental procedures were approved by the Medical Experimental Animal Administrative Committee of Fudan University. Mice were housed in specific pathogen free (SPF) conditions in ventilated caging system with a density lower than 0.01m^2^ per animal. The circadian were set at 8 a.m. in a 12h/12h light/dark cycle with temperature at 22 ± 2 °C and 45 - 65% relative humidity. Mice have access to sterilized water and irradiated phytoestrogen-free standard diet for rodents (Beijing Keao Xieli Feed Co., LTD.) ad libitum.

APP/PS1 ((Jax stock #34829) mice and Ai9 (RCL-tdT) (Jax stock #007909) mice were purchased from the Jackson Laboratory. *Tpk^foxedl/foxed^* mice were generated by our lab according to the description in the paper accompanied and published before ^1^. *Cx3cr1^CreERT2/+^* transgenic mice were obtained from Prof. Yuqiu Zhang at the Institutes of Brain Science, Fudan university, which are available from the Jackson Laboratory. Microglia-specific *Tpk* knockout *Cx3cr1^CreERT2/+^; Tpk^foxedl/foxed^* mice (mcKO mice) were generated by crossing *Tpk^foxedl/foxed^* mice with *Cx3cr1^CreERT2/+^* mice. We next crossed mcKO mice with the *Ai9* or the *APP/PS1* mouse strains to generate *Ai9/+; Cx3cr1^CreERT2/+^; Tpk^foxed/foxed^* mice (mcKO mice with tdTomoto signaling in microglia) or *APP/PS1; Cx3cr1^CreERT2/+^; Tpk^foxed/foxed^* mice (APP/PS1/mcKO mice) respectively. To conditionally knockout *Tpk* gene in microglia of all types of transgenic mice, each animal at 12 weeks of age were intraperitoneally injected with tamoxifen (50 mg / kg·day) for five consecutive days.

3. **Cell lines culture**

All culture medium and supplements were purchased from Gibco, USA except otherwise informed. Cell lines were purchased from Fuheng Biology Science and technology Co., Ltd (Shanghai, China) with report of cell line identification. Mouse BV2 microglia and human HMC3 microglia were maintained in high glucose Dulbecco’s modified Eagle’s medium (DMEM) supplemented with 10 % fetal bovine serum (FBS) and 100 mg / mL penicillin/streptomycin (PS) under humidified 5 % CO2 / 95 % air environment at 37℃. L929 cell line was used for preparing conditioned medium (LCCM). L929 cell line were maintained under the same condition and supplements but DMEM/F12 medium. The supernatant of L929 culture was collected by discarding precipitation after centrifuge at 300 x g for 10 minutes 2 days after 90% confluency met by the cultured cell. Then the LCCM was preserved at -80°C shortly before use.

4. **Primary microglia culture**

Primary microglia were extracted and cultured according to Gao et al^2^. Brain from neonatal mice were obtained and immersed in ice-cold HBSS with 1M HEPES. HBSS washed tissue were mechanically dissected into pieces less than 1 mm^3^ after removal of pia mater under stereoscope (Nikon, Japan). Then all tissue were collected by centrifuge at 1,000 x g and transferred into DMEM with 0.25% trypsin. After incubated at 37°C for 30min with gentle shaking at 5 minutes interval, same amount of DMEM with 10% FBS was added to terminate digestion. DNase I (Invitrogen, NY, USA) was then applied to yield single cell suspension. The yield was centrifuged at 300 x g for 5 min. Then the precipitant was resuspended and seeded into poly-D-Lysine (Sigma, P7280) treated T75 flasks (Thermo Fisher Scientific, USA) by DMEM/F12 containing 10% FBS and 1% PS, which was incubated at 37°C with 5% CO_2_ for 8 days with change of the medium at an interval of 4 days. At 90% confluency met by the mixed glial cells, L929 cell line conditioned medium (30% v/v) was added into the DMEM/F12 + 10% FBS + 1% PS medium to favor microglia proliferation. After 2 days incubation, microglia was detached by shaking the flasks at 150 rpm 37°C for 1 hour. The supernatants were collected and flited by 40 µm filters (Falcon, USA). Cell pellet collected by centrifugation at 300 x g for 5 min was then resuspended in DMEM/F12 medium with 30% LCCM and seeded on a PDL-treated 6-well cell culture dish at the density of 1×10^6^ cells per well for subsequent experiments.

5. **Ex Vivo microglial TPK knockout**

For ex vivo *Tpk* knockout model, primary microglia seeded in 6-well dish were kept in standard condition with 12-24 h adherence time reserved. Primary microglia with normal morphology were treated by 1 µM 4-hydroxytamoxifen or PBS for 5 consecutive days. Then the expression level of TPK was confirmed by western blotting and those batch with more than 60% down regulation were selected for subsequent experiments.

6. **In vitro LPS treatment**

For simulating acute inflammatory reaction *in vitro*, 500 ng / mL of LPS or vehicle control (PBS) were added into the culture medium of LPS group or control group respectively, and the cultured cells were harvested for analysis at 24 hours after treatment.

7. ***In vivo* LPS treatment**

For simulating acute inflammatory reaction in vivo, three-month-old mice were randomly assigned to treatment groups and were given a single intraperitoneal injection of lipopolysaccharides (LPS) at 5 mg per kg bodyweight (L7261, Sigma, USA) or vehicle injections (PBS) at 24 hours before taking brain samples.

8. **Microglia isolation from adult mice**

The adult mouse microglia were isolated as previously described ^3^. Mice were anesthetized by sodium pentobarbital before perfused with ice-cold phosphate buffered solution (PBS). After removal, the brains were immediately immersed in PBS solution consisting Liberase TL (Roche, Germany), DNase I (Invitrogen, USA), and RNase inhibitor (Promeger, USA), and mechanically separated with microscopic scalpel and forceps. The separated brain tissues were then incubated in the mixed enzyme solution at 37 °C for 30 minutes. Then, 20 μL (0.5 mM) of ethylenediamine tetraacetic acid (EDTA) were added to terminate the enzymatic reaction. The solution was pipetted repeatedly and filtered with 70 μm cell strainer (Falcon, Germany) to obtain single-cell suspension. After centrifugation at 300g and 4 °C for 10 minutes, the supernatant was discarded. Then, myelin and debris in the single-cell suspension were removed by applying myelin removal MicroBeads (Miltenyi Biotec, 130-109-398) and LD column (Miltenyi Biotec, 130-042-901). The remaining cell suspension was further sorted for subsequent experiments. For Bulk RNA-seq and single-step RT-PCR assay, cells were stained with anti-CD11b (Biolegend, 101205), anti-CD45 (Biolegend, 103113) and APC anti-Ly6c (Biolegend, 128015). The CD11b^+^CD45^lo^Ly6c^-^ cells in the brain were fluorescence-activated cell-sorted (FACS) directly into lysis buffer. For immunoblotting experiments, the cell suspension was incubated with CD11b MicroBeads (Miltenyi Biotec, 130-049-601) at 4°C for 15 minutes and centrifuged at 300g, 4°C for 10 minutes. The pellets were collected, resuspended, and passed through LS column (Miltenyi Biotec, 130-042-401) to perform the positive classification sorting. The solution containing CD11b^+^ cells were centrifuged at 500g, 4 °C for 10 minutes. The pellets were collected in radioimmunoprecipitation assay (RIPA) lysis buffer containing protease inhibitor. All samples were frozen on dry ice and stored at -80 °C until further processing.

9. **In vitro treatment of A**β

The synthetic Aβ peptide was purchased from Bachem (Bubendorf, Swizerland). According to manufacturer’s protocol, both Hexafluoroisopropanol (HIFP) pre-treated Aβ1-42 (4090148) and Aβ42-1 (4107743) peptides were suspended in Dimethyl sulfoxide (DMSO) at 5 mM concentration and sonicated 20 s for full dissolve. The solution was diluted by corresponding medium for each type of cell culture. The peptide-containing medium were changed into 90% confluence cultures and incubated in humidified 5 % CO_2_ / 95 % air environment at 37 °C. After 12 h incubation, cells were digested from petri dish by 0.25% trypsin and collected by centrifuge at 300 x g for 10 minutes. The cell pellets were treated with RIPA lysis buffer and preserved in -80 °C before use.

10. **Phagocytosis assay**

Incubate FluoSpheres^TM^ carboxylate (Invitrogen, USA), with 10% FBS in PBS at 37 °C to block surface antigen on the beads. The solution was then centrifuged at 4,000g for 20 min. The pellet was resuspended with culturing medium and added into both groups of wild type and TPK knockout primary microglia culture. After 1 h incubation at 37 °C, the cultures were washed with PBS for 3 times to remove free floating beads in the medium. The attached microglia were fixed by 4% fromalinhyde (PFA). Fluorescent signal was observed under fluorescence microscope (Nikon).

11. **Microglia SMART-seq2**

For both mcKO and littermate mice, 500 FACS-sorted CD11b^+^CD45^lo^Ly6c^-^ microglia cells from each brain were directly dropped into lysis solution including triton. Then, the SMART-seq2 for microglia were performed by 10K Genomics, Shanghai, China. Briefly, the process involves the following steps: Reverse transcription of RNA into cDNA; Amplify and purify the cDNA; Fragment cDNA with Tn5 enzyme and construct the library. Firstly, the lysate samples were vortexed, centrifuged, and placed on a preheated PCR machine for heat-treatment. Then, Smart RT buffer and RT enzyme were added, and the reverse transcription reaction was carried out according to the program. Then, Smart Amp buffer-1 and Smart Amp enzyme-1 were added to the cDNA product obtained in the previous step and the amplification reaction was carried out according to the procedure. A small amount of cDNA product was token out for quality control by running electrophoresis gel and quantitative purification of cDNA was performed using a DNA purification kit. The purified cDNA samples were added to TN5 buffer and TN5 enzyme-1, and the reaction was interrupted according to the program. After the reaction, Smart Amp buffer-2, Smart Amp enzyme-2, S5XX, N7XX was mixed for library construction, purify, and send for testing. Illumina NovaSeq 6000 sequencing platform was used for sequencing and PE150 sequencing mode was adopted.

12. **SMART-seq2 data analyses**

The SMART-seq2 data analyses were also performed by 10 K Genomics, Shanghai, China. Briefly, FastQC software was used to evaluate the quality of the original data and HISAT2 software was adopted to compare the filtered data to the reference genome. Then Stringtie was used to calculate the gene expression and normalize it to generate the gene expression normalization file required for subsequent analysis. Gene expression standard files were then used for further principal component analysis of samples, and software such as edgeR and DEseq2 were used to screen differentially expressed genes between groups, then GO and KEGG enrichment analysis of differential genes was performed.

13. **single-step RT-PCR**

Single-step RT-PCR for isolated microglia from adult mice brain was performed according to Yazawa et al^4^. Microglia isolated from FACS (500 cells) was instilled into 11 μL ice-cold master mix of Cells Direct one-step for qRT-PCR (Invitrogen, 46-7200), composed of 10 μL resuspension buffer and 1 μL lysis enhancer. After thorough mixing, the sample was frozen on dry ice and stored at -80°C before further process. When in use, the samples were thawed at 4 °C and sat in 75 °C for 10 minutes, before put in 4 °C to refrigerate for 5 minutes. According to manufacturers’ protocol, add 10 μL reaction buffer mix and 2 μL RT ENZYME mix from Superscript III first-strand synthesis supermix for qRT-PCR (Invitrogen, 11752-050). The samples were mixed well by vertexing for 10s, and went through the reverse transcription procedure: 25 °C for 10 minutes, 50 °C for 30min, 85 °C for 5 minutes and preserved at 4 °C. Then, 1 μL RNase H was added and mixed at 37°C for 20 minutes to eliminate all remaining RNA.

14. **qRT-PCR**

Total RNA was extracted from cell sample or brain tissues of mice using the TRIzol reagent (Invitrogen). For Real-time RT-PCR experiments, complimentary cDNA was synthesized using the PrimeScript™ RT reagent Kit with gDNA Eraser (Takara, Japan) according to the manufacturer's protocol. Primers were designed using online Primer-BLAST software according to the instructions. The cDNA synthesized by RNA sequencing was also verified by Real-time PCR. Real-time PCR was performed using SYBR Green Master Mix (TaKaRa, Japan) on 7500 Real-time PCR system (Applied biosystems). All reactions were performed in triplicates, and the results normalized to that of control groups from the same preparation.

15. **ELISA**

The levels of Aβ40 and Aβ42 were determined by ELISA (KHB3441, Invitrogen, USA) according to the manufacturer’s instructions. Brain tissues were weighed and homogenized in ice-cold PBS containing the protease inhibitor cocktail (Complete Protease Inhibitor Cocktail, Roche Diagnostics) followed by centrifugation for 20 min at 15,000 x g, 4°C, guanidine buffer (5 M guanidine HCl/50 mM Tris-HCl, pH 8.0). The homogenates were mixed for 4 hours at the room temperature and were diluted 1: 5 in PBS containing 5% BSA and 0.03% Tween-20 supplemented with the protease inhibitor cocktail followed by centrifugation at 16,000 x g for 20 min at 4°C. The supernatant was diluted and analyzed according to the manufacturer’s instructions.

16. **Western blotting**

Cell samples were resuspended in radioimmunoprecipitation assay (RIPA) lysis buffer (1% Triton X-100, 1% sodium deoxycholate, 0.2% sodium dodecyl sulfate (SDS), 0.15 M NaCl, 0.05 M Tris-HCl, pH 7.2) containing the protease inhibitor mixture and PhosSTOP phosphatase inhibitor cocktail (Roche). Brain tissues were weighed and lysed by the RIPA lysis buffer at 0.01 mg / μL. Protein concentrations were determined with the Pierce™ BCA protein assay kit according to the manufacturer’s instruction (Thermo Scientific, Rockford, IL). The same amount of proteins was loaded in each lane for electrophoresis on a 10% denaturing Tris/glycine sodium dodecyl sulfate (SDS) polyacrylamide gel and subject to electrophoresis. Proteins were transferred to the polyvinylidene fluoride membranes (Millipore) and blocked in 5% milk in Tris-buffered saline supplemented with 0.1% Tween-20 (TBS-T, pH 7.4) for 1 h. The membrane was incubated overnight at 4^o^C with TBST containing 5% milk and primary antibodies. Then, membranes were washed in TBS-T and incubated for 1 h with horseradish peroxidase (HRP)-conjugated anti-mouse or anti-rabbit IgG (Millipore) in TBS-T containing 5% milk. After a final wash in TBST, membranes were incubated with ECL substrate (Pierce® Fast Western Blot Kit, 35050, USA) for 1 - 2 minutes, signals were detected with Tanon 6200 Luminescent Imaging Workstation (Tanon Science & Technology Co., Shanghai, China). Results were quantified using Image J (N.I.H.).

17. **Immunohistochemical staining**

After deeply anesthesia, mice were intracardially perfused with PBS and then 4% paraformaldehyde for fixation. Serial coronal sections (25 μm) were cut with a sliding microtome (Leica) and stained using a freely floating method. After being washed in PBS at pH 7.4, the sections were put into PBS containing 5% bovine serum albumin (BSA) and 0.5% Triton X-100 for 2 h at 37 °C for 1 h in order to block non-specific reactions. Then, the sections were incubated with primary antibodies overnight at 4 °C. After being washed with PBS for 3 times, 10 minutes each time, the sections were incubated with corresponding secondary antibody conjugated to Alexa Fluor 488 (Invitrogen, 1:500), Alexa Fluor 555 (Invitrogen, 1:500), or Alexa Fluor 647 (Invitrogen, 1:500), for 2 h at 37 °C. For examination of amyloid plaques, the sections were incubated in 0.02% Thioflavin S at the room temperature (22-25 °C) for 30 min. Nuclei were stained with DAPI (Sigma, D9542, 1:1000) and the slices mounted on 3-aminopropyltriethoxysilane (APES)-coated glass slides. Z-stack images were taken using Nikon A1 (Tokyo, Japan) laser scanning microscope, with 20-x, 40-x, or 60-x objectives. Results were quantified using ImageProPlus (Media Cybernetics, Silver Spring, MD, USA).

18. **Electron microscope**

CKO mice and their littermates (n = 4 per group) were anesthetized and transcardially perfused with 0.1 M phosphate buffer (PB, pH 7.4). Brains were removed and approximately 1 cm^3^ cortical tissue at the same anatomical location were extracted and fixed into 2.5% glutaraldehyde in 0.1 M phosphate buffer overnight at 4 °C. Then these tissues were rinsed with 0.1 M PB, postfixed in 1% Osmium (Electron Microscopy Sciences) for 2 hours, rinsed with 0.1 M PB, dehydrated in a graded series of ethanol and embedded. Ultrathin sections (about 100nm) of each tissue were cut using a LEICA EM UC7 Ultramicrotome (Leica) and stained with lead citrate (3%), and observed by using a H-7650 transmission electron microscope (Hitachi). As descripted in the literature, microglia were identified using a combination of ultrastructural characteristics, including highly electron-dense cytoplasm and nucleus, irregularly shaped nucleus, and a cytoplasm rich in free ribosomes and vesicles ^5^. Ten ultrathin sections were analyzed per mouse in order to evaluate the ultrastructure of microglia.

19. **Mass cytometry**

Brain tissue was taken from mcKO mice and their littermates (n = 6 per group) after transcardially perfusion unconsciously. The tissue samples were lysed in 1 % SDS with 8M urea and protease inhibitor cocktail with incubation at 4 °C for 30 min with shake at 10 minutes interval. Un-lysed matter was discarded after centrifugation at 12000g at 4 °C for 20min. The protein concentration was determined by BCA Protein Assay Kit. For digestion, 100 μg protein in 100 μL lysis buffer from each sample was reduced with 2 μL 0.5 M Tris-phosphine (TCEP) at 37 °C for 60min. Then, 4 μL of 1 M iodoacetamide (IAM) were added to alkylate the protein at room temperature for 40 minutes in darkness.

To precipitate the protein, five volumes of cold acetone were added and incubated at -20 °C overnight. After centrifugation at 12000 g at 4 °C for 20 minutes, the pellet was washed twice by 1 mL pre-chilled 90% acetone aqueous solution and re-suspended with 100 μl 10 mM riethylammonium bicarbonate (TEAB) buffer. For peptide generation, 1 : 50 (m:m) trypsin (Promega, Madison, WI) was added and incubated at 37 °C overnight. The digested peptide mixture was then desalted by C18 ZipTip, and quantified by PierceTM Quantitative Colorimetric Peptide Assay (23275) before lyophilized by SpeedVac. Re-dissovle the samples in 20 mM ammonium formate, pH 10.0 before fractionating by high pH separation using Ultimate 3000 system (ThermoFisher scientific, MA, USA) connected to a reverse phase column (XBridge C18 column, 4.6 mm x 250 mm, 5 μm, (Waters Corporation, MA, USA). After re-equibrating for 15 minutes and flow at 1 mL/min, 30°C, 12 fractions were collected and dried in a vacuum concentrator. The collected fractions were analyzed by online nano flow liquid chromatography tandem mass spectrometry performed on an EASY-nanoLC 1200 system (Thermo Fisher Scientific, MA, USA) connected to a Orbitrap FusionTM LumosTM TribridTM mass spectrometer (Thermo Fisher Scientific, MA, USA). Raw Data collected by data dependent acquisition of the spectrometer were processed and analyzed by Spectronaut 15 (Biognosys AG, Switzerland). Spectronaut was set up to search the database of uniprot_Mus_musculus (version 201907, 22290 entries) database. Since the database was established, the mass spectrometer was then run under data independent acquisition (DIA) mode. Raw Data of DIA were processed and analyzed by Spectronaut 15 (Biognosys AG, Switzerland). Data extraction was determined by Spectronaut 15 based on the extensive mass calibration. Qvalue (FDR) cutoff on precursor level was 1% and protein level was 1%. After Welch's ANOVA Test, different expressed proteins with p value <0.05 and |fold change| >1.2 were filtered.

20. **In vivo imaging of microglia with two-photon microscopy.**

The mcKO mice with tdTomoto signaling in microglia and their littermate control (HET and WT) were used for in vivo imaging of microglia with two-photon microscopy. Surgical procedures were performed as previously described ^6^. Briefly, adult mice (5-12 months old) were anesthetized with isoflurane (4–5% for induction; 1–1.5% for maintenance) and body temperature was kept at 36-37°C. Dexamethasone (2 mg per kg) and carprofen (5 mg per kg) were given subcutaneously. Hair, skin and periosteum overlying the neocortex were removed. A 4 mm diameter circle was drilled over somatosensory cortex, then both the skull piece and the dura underneath were carefully removed by fine forceps. Sterile saline was frequently dripped on the surgical area to avoid overheating and keep tissue moist. A 4 mm cover glass was gently pressed onto the brain surface and glued to the skull. A customized head bar was glued onto the skull. Mice were placed onto a heating pad for recovery. Head-fixed imaging procedures were operated only after mice were widely awake, that is approximately 2 hours after the surgery.

Imaging was performed using a two-photon microscopy equipped with a Ti-sapphire tunable laser (Thorlabs), a gallium arsenide phosphide (GaAsP) detector (Thorlabs) and a ×16/0.8 NA water-immersion objective (Nikon). The tdTomato labelled microglia were imaged through excitation at a wavelength of 1000 nm. Average laser power was <10–30 mW at the tissue surface and adjusted with depth exponentially as needed to compensate for signal loss due to scattering and absorption. First, we recorded a Z and T series base activity of microglia in layers 1 and 2 of the somatosensory cortex. Z-stacks were acquired at 35-μm axial step size, and used a three-frame average, 1024 × 1024-pixel resolution and ×2.0 zoom. Recordings lasted for about 30 minutes.

Then a longer (about 45 minutes) Z and T series was recorded after laser ablation was made using the same parameters described above. Laser-induced focal ablation is a highly reproducible injury model to observe the response of microglia^7^. To execute a laser ablation, wavelength of the two-photon laser was tuned to 800 nm and the laser power was 50–60 mW at the sample., Z position was adjusted to the center of Z stacks, and region of interest was adjuster to 2048 × 2048-pixel resolution and ×37.6 zoom. Just 1 - 3 seconds of exposure to the laser induced a focal injury site. Recording was started immediately after the laser ablation. For analyses of two-photon image stacks of baseline motility, we used a customized MATLAB script that makes adaptive threshold in selected regions of interest to automatically generates a mask for the microglia cell and processes at each time point. The motility was calculated as the differences in the cell masks between two timepoints, normalized by the total cell area. For the tissue ablation experiment, we measured the amount of all processes within 20 microns radius from the ablation site.

21. **Quantification of microglia barrier**

Quantification of microglia coverage of plaques were quantified following previously described methods ^8^ using a customed MATLAB script. Briefly, all image files from various groups were processed together in a randomization and blinding process to mask the file names. Then the blinded stack was projected at the center of the plaque. Microglia processes were detected by a global threshold to the image, and the overlap between the microglia mask and plaque mask were accounted as microglia coverage. The coverage was then transformed into angular measurements for normalization. Users were allowed to manually adjust for the coverage during this process. After all image files were processed, the files will then be unblinded and group information revealed.

22. **Statistical analysis**

GraphPad Prism 8 (version 8.4.3 (471), GraphPad software) was used for statistical analyses. Student’s t-test for single comparisons or one-way ANOVA for multiple comparisons with appropriate Tukey’s or Dunnett’s Multiple Comparison tests were used to determine statistical differences. Summary results are shown as means ± SEM.
