## Supplementary figures and table for "The mechanism and consequences of amyloid-β modulating thiamine pyrophosphokinase-1 expression in microglia"

**Supplementary Figures and Tables:**

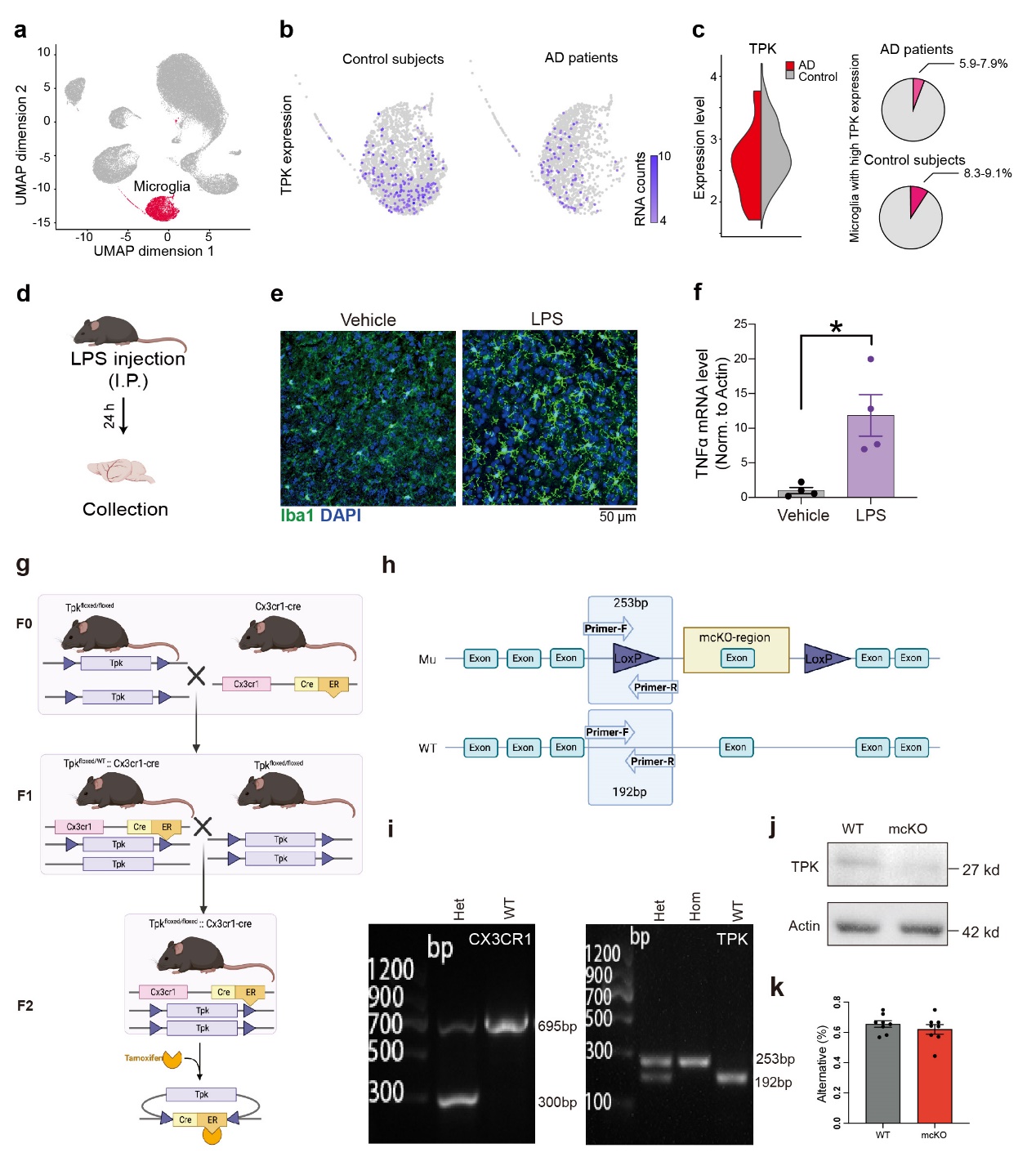

**Extended figure 1. Human brain single-cell sequencing data and the generation and validation of mouse models.**

**a-c**, Human brain single-cell sequencing data reanalysis from a previously published single-cell transcriptome dataset from 24 human AD and 24 control subjects. **a,** 2-dimensional Uniform Manifold Approximation and Projection (UMAP) projection of all annotated cells (n=75,060). **b,** *Tpk* mRNA expression in microglia clusters of 24 AD patients and 24 control subjects. **c,** Quantification of *Tpk* mRNA expression level in AD patients and control subjects. Violin plots (left panel) showed AD patients had a lower *Tpk* mRNA level as compared with control subjects. Pie plots (right panel) showed the proportion of microglia with high TPK content in AD patients is higher than that in the control subjects.

**d,** The strategy for constructing an acute inflammatory mice model by intraperitoneal injection of LPS at 5 mg/kg and sampling the brain 24 h after treatment.

**e, f,** Confirmation of establishment acute inflammatory model successfully by observing Iba1 immunolabeled microglia in cortical of mice injected with 5 mg/kg LPS vs. vehicle. Representative confocal images showed significantly changed morphology with features representing activation, like enlarged cell body and ramified processes (**e**). The relative mRNA level of brain TNFα detected by real-time PCR in each group was shown elevated mRNA level of TNFα in LPS stimulated mice compared to vehicle also indicates inflammatory activation (**f**). n = 4 mice per group.

**g**, The schematic illustration of crossbreed strategy. The homozygotic mcKO mice is breeding *Tpk^floxed/floxed^* and *Cx3cr1^CreERT2/+^* mice for two generations. Genotypes were annotated under each mouse, with purple triangle indicates FloxP site.

**h,** The gene structure of *Tpk^floxed/floxed^* (upper panel) and wild type (lower panel) mice with genotyping primers’ annealing sites indicated. The light blue rectangles indicate 5 exons in the TPK gene. The LoxP sites were inserted at ends of the 4^th^ exon, constraining the mcKO region. The forward and revers primers were designed to anneal at upper and lower stream of the first LoxP site, which will generate 253 bp or 192 bp product in *Tpk^floxed/floxed^* and wild type mice respectively. **g,** Representative image of genotyping results of *Cx3cr1^CreERT2/+^* (left panel) and *Tpk^floxed/floxed^* (right panel) mice. For *Cx3cr1^CreERT2/+^*, wild type mice have one band at 695 bp and heterozygotic mice have an extra band at 300 bp in the PCR gel electrophoresis.

**i,** Confirmation of *Tpk* knockout detected by western blotting. Microglia isolated from tamoxifen induced mice showed significant reduction of TPK.

**j**, Confirmation of TPK protein expression in isolated microglia of mcKO mice and their littermates 3 month after tamoxifen induction. (n = 3 : 3)

**k**, Y maze test showed no significant difference between mcKO and their littermates. (n = 8 : 8).

Statistical analyses were performed using unpaired two-tailed t-tests. Summary data represent means ± SEM. Animal number per group is listed in statistical graphs. * p < 0.05. Scale bars, 50 μm (**e**).

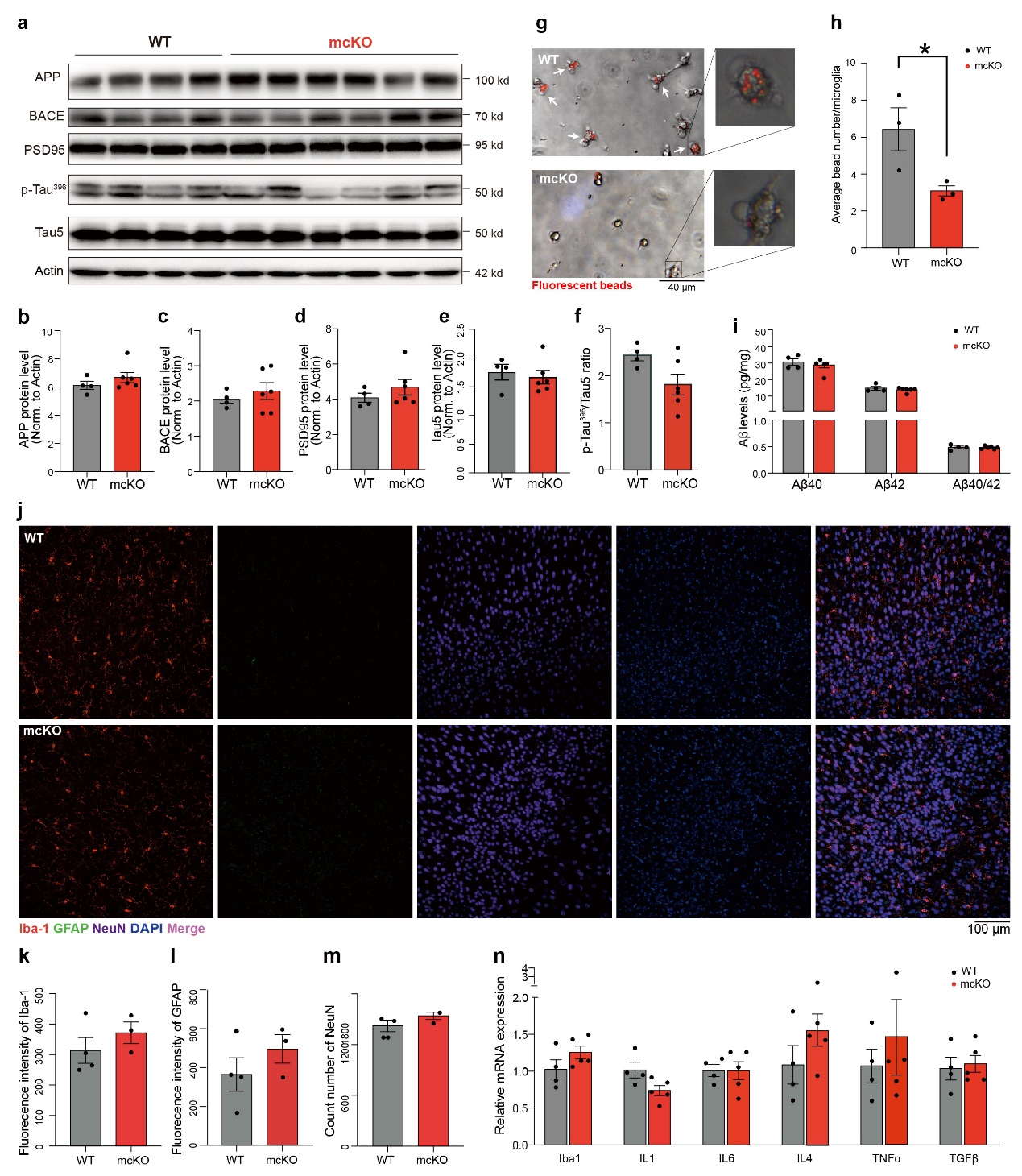

**Extended figure 2. Additional measurements of mcKO mice.**

**a-f,** Representative western blotting images (**a**) and quantification of protein levels in brain tissue of 18-month-old mcKO mice and their littermates (n = 6 : 4). No significant difference was found in APP (**b**), BACE (**c**), PSD95 (**d**)**,** Tau5 (e) or p-Tau^396^/Tau5 ratio (**f**) between mcKO mice and their littermates.

**g, h.** Phagocytosis was measured in primary microglia of mcKO or control mice treated with fluorescent beads. Representative phagocytosis images were shown in **g** with ROI zoomed in showing multiple individual beads engulfed by microglia (right panel). Arrows indicate microglia that engulfed with beads, n = 3 independent experiments. Quantification of the average beads numbers per microglia was significantly decreased in mcKO primary microglia as compared with that in control primary microglia (**h**).

**i.** Quantification of the protein level of Aβ 40, Aβ42 and their ratio detected by ELISA in brain tissue of 6-month-old mcKO mice and their littermates (n = 4 : 6). No significant difference was found in either proteins or Aβ40/42 ratio between the two group.

**j-m,** representative image (**j**) and quantification of immunofluorescent staining of Iba1 (red), GFAP (green), NeuN (purple) and DAPI (blue). The fluorescence intensity of Iba1 (**k**) or GFAP (**l**) showed no significant difference between control and mcKO. The count number of NeuN was similar between two groups (**m**). (n = 4 : 3)

**n,** The transcription level of Iba1 and selected inflammatory factors. There was no significant difference between control and mcKO mice (n = 4 : 5).

Statistical analyses were performed using unpaired two-tailed t-tests. Summary data represent means ± SEM. Animal number per group is listed in statistical graphs. * p < 0.05. Scale bars, 40 μm (g, left), 100 μm (**j**).

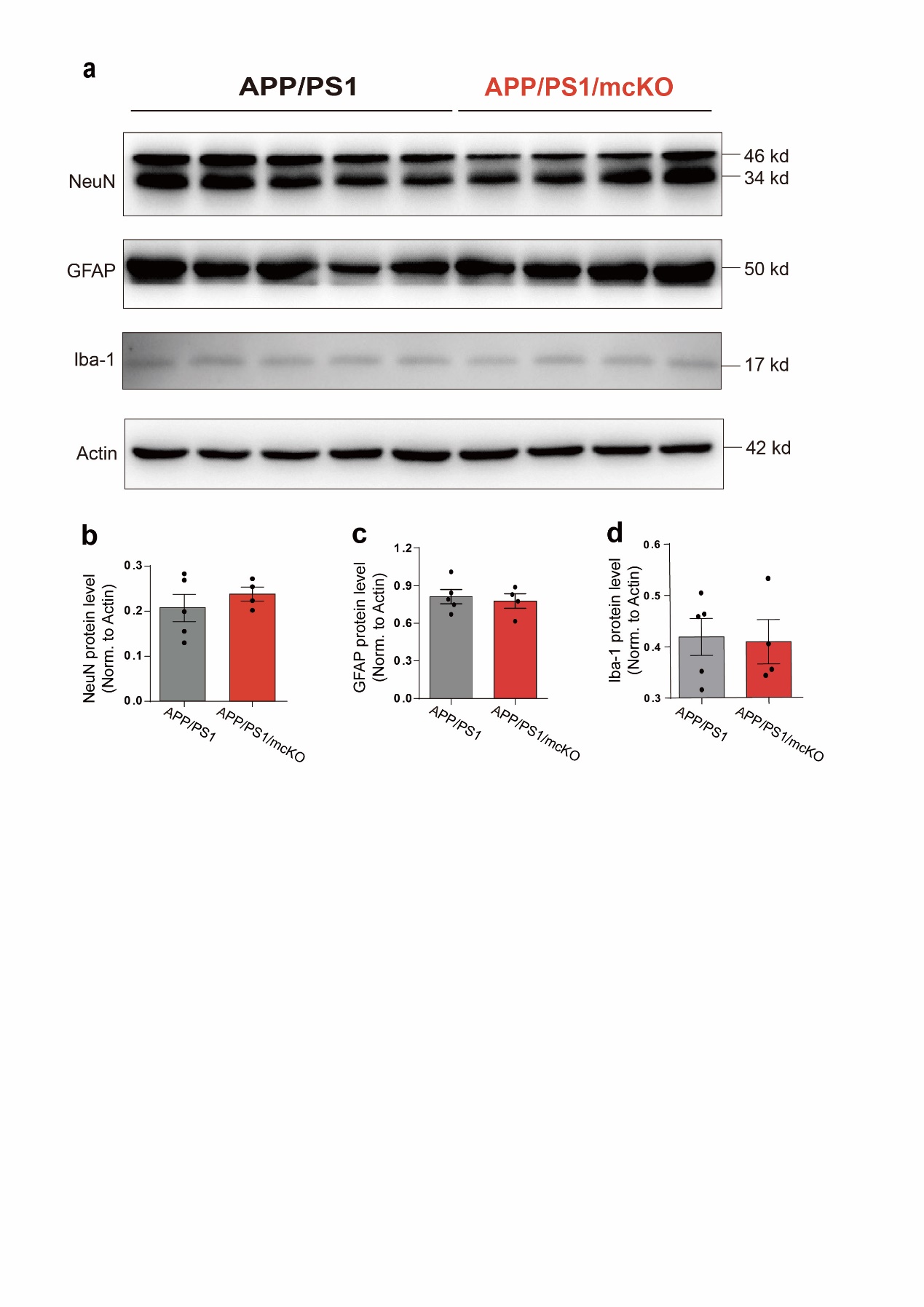

**Extended figure 3. Additional measurements of APP/PS1/mcKO mice.**

**a-d**, Representative western blotting images (**a**) and quantification of protein levels in brain tissue of 18-month-old mcKO mice and their littermates (n = 4 : 5). No significant difference was found in NeuN **(b)**, GFAP **(c)** or Iba-1 **(d)** between APP/PS1/mcKO mice and their APP/PS1 littermates.

Statistical analyses were performed using unpaired two-tailed t-tests. Summary data represent means ± SEM. Animal number per group is listed in statistical graphs. * p < 0.05.

| **Gene_Name** | **log2**  **FoldChange** | **stat** | **pvalue** | **padj** | **CKO1** | **CKO2** | **CKO3** | **WT1** | **WT2** |
| --- | --- | --- | --- | --- | --- | --- | --- | --- | --- |
| Hsp90aa1 | -3.67925 | -11.7548 | 6.67E-32 | 1.01E-27 | 147.2165 | 102.7132 | 109.9412 | 1758.328 | 1317.298 |
| Csf1 | -4.55099 | -10.607 | 2.76E-26 | 2.09E-22 | 6.110875 | 11.82758 | 14.21009 | 229.277 | 265.6552 |
| Fhad1 | -3.70623 | -9.94241 | 2.72E-23 | 1.37E-19 | 42.22059 | 41.70779 | 35.89917 | 692.6749 | 349.0841 |
| Gem | -3.54164 | -9.91223 | 3.68E-23 | 1.39E-19 | 25.55457 | 24.27767 | 35.15127 | 310.0083 | 346.8886 |
| Slc15a3 | -3.60022 | -9.815 | 9.70E-23 | 2.94E-19 | 71.6639 | 65.98546 | 82.26893 | 565.1194 | 1214.11 |
| Jun | -2.54398 | -9.10363 | 8.74E-20 | 2.20E-16 | 463.3154 | 369.1451 | 379.9329 | 2363.814 | 2351.378 |
| Ppp1r15a | -4.41331 | -8.83163 | 1.03E-18 | 2.23E-15 | 13.88835 | 3.735026 | 13.46219 | 150.1603 | 292.0012 |
| Nfkbiz | -2.75769 | -8.7285 | 2.58E-18 | 4.88E-15 | 165.5492 | 111.4283 | 113.6807 | 837.9913 | 926.4999 |
| Rasgef1b | -2.57819 | -8.60532 | 7.61E-18 | 1.28E-14 | 104.4404 | 117.0308 | 130.8824 | 674.914 | 726.7096 |
| Chka | -3.18515 | -8.56093 | 1.12E-17 | 1.69E-14 | 53.88681 | 72.2105 | 62.07565 | 374.5934 | 768.4241 |
| Gcnt1 | -2.36388 | -8.16429 | 3.23E-16 | 4.45E-13 | 185.5484 | 183.0163 | 192.2101 | 860.596 | 1064.816 |
| Zbtb2 | -3.01556 | -8.12303 | 4.55E-16 | 5.74E-13 | 48.33147 | 39.21777 | 64.31935 | 306.7791 | 511.5509 |
| Bcl6 | -3.3663 | -8.05027 | 8.26E-16 | 9.62E-13 | 23.33243 | 11.20508 | 21.68908 | 195.3698 | 191.0083 |
| Casp4 | -2.94117 | -7.93902 | 2.04E-15 | 2.20E-12 | 24.99903 | 32.99273 | 27.67228 | 235.7355 | 201.9858 |
| Glul | -2.81834 | -7.89886 | 2.81E-15 | 2.84E-12 | 243.8795 | 112.6733 | 154.8152 | 1323.994 | 1082.38 |
| Atf3 | -2.74083 | -7.87668 | 3.36E-15 | 3.18E-12 | 192.7703 | 111.4283 | 116.6723 | 773.4062 | 1106.531 |
| Hsp90ab1 | -2.42271 | -7.86441 | 3.71E-15 | 3.30E-12 | 398.318 | 316.2322 | 286.4455 | 2100.629 | 1477.57 |
| Malat1 | -2.43248 | -7.78001 | 7.25E-15 | 6.10E-12 | 1450.5 | 817.9707 | 1210.849 | 6282.513 | 6239.604 |
| Gm15459 | -3.21975 | -7.54587 | 4.49E-14 | 3.58E-11 | 36.66525 | 23.65517 | 20.19328 | 334.2277 | 166.8578 |
| Junb | -3.08729 | -7.52864 | 5.13E-14 | 3.88E-11 | 32.22098 | 19.92014 | 26.17648 | 282.5597 | 160.2713 |
| Tent5c | -2.28836 | -7.45158 | 9.22E-14 | 6.65E-11 | 127.7728 | 98.97819 | 112.1849 | 574.8071 | 529.1149 |
| Klf6 | -2.71863 | -7.42852 | 1.10E-13 | 7.56E-11 | 173.8822 | 93.37565 | 88.25213 | 892.8886 | 669.6267 |
| Gm34455 | -2.79256 | -7.37017 | 1.70E-13 | 1.12E-10 | 78.33031 | 77.81304 | 43.37816 | 578.0364 | 344.6931 |
| Runx1 | -2.30548 | -7.28428 | 3.23E-13 | 2.04E-10 | 279.4337 | 220.989 | 317.8572 | 1088.258 | 1609.3 |
| Pde3b | -2.40549 | -7.25762 | 3.94E-13 | 2.39E-10 | 408.8731 | 288.842 | 569.1514 | 2178.131 | 2294.295 |
| Dennd4a | -2.03729 | -7.20814 | 5.67E-13 | 3.30E-10 | 359.4306 | 332.4173 | 407.6052 | 1401.496 | 1607.104 |
| Msn | -2.31057 | -7.17319 | 7.33E-13 | 3.98E-10 | 80.55245 | 64.11795 | 80.02523 | 369.7495 | 373.2346 |
| Arhgef1 | -2.24412 | -7.17259 | 7.36E-13 | 3.98E-10 | 78.33031 | 82.79308 | 79.27733 | 379.4373 | 379.821 |
| 1700001K19Rik | -3.49587 | -7.14254 | 9.16E-13 | 4.78E-10 | 6.666409 | 19.92014 | 15.70589 | 200.2137 | 114.1659 |
| Smad7 | -2.76387 | -7.10688 | 1.19E-12 | 5.87E-10 | 38.88739 | 96.48817 | 62.07565 | 427.8761 | 465.4454 |
| Neat1 | -2.49087 | -7.10518 | 1.20E-12 | 5.87E-10 | 86.10779 | 71.588 | 89.00003 | 335.8424 | 590.5888 |
| Ifrd1 | -2.89988 | -7.09353 | 1.31E-12 | 6.05E-10 | 144.9944 | 65.98546 | 86.00843 | 481.1588 | 1001.147 |
| Mef2d | -2.51123 | -7.09221 | 1.32E-12 | 6.05E-10 | 69.44176 | 46.06532 | 84.51263 | 368.1349 | 390.7985 |
| Adap2 | -2.2681 | -7.07815 | 1.46E-12 | 6.50E-10 | 124.9952 | 94.62066 | 124.1513 | 628.0898 | 474.2274 |
| Zfand5 | -2.61841 | -7.0563 | 1.71E-12 | 7.40E-10 | 61.10875 | 37.97277 | 58.33615 | 255.111 | 390.7985 |
| Cntnap5a | 2.963722 | 6.985655 | 2.84E-12 | 1.19E-09 | 185.5484 | 307.5171 | 246.0589 | 32.29253 | 30.73696 |
| Sox4 | -3.50759 | -6.95103 | 3.63E-12 | 1.48E-09 | 30.55438 | 12.45009 | 46.36976 | 492.4611 | 180.0308 |
| Hspa1a | -3.40323 | -6.94537 | 3.77E-12 | 1.49E-09 | 44.99826 | 11.20508 | 35.15127 | 432.72 | 210.7677 |
| Fosb | -2.65376 | -6.94256 | 3.85E-12 | 1.49E-09 | 123.8841 | 84.66059 | 49.36136 | 526.3683 | 559.8518 |
| Tmx4 | -2.57485 | -6.85742 | 7.01E-12 | 2.62E-09 | 194.9925 | 113.2958 | 81.52103 | 724.9674 | 827.7025 |
| Etf1 | -2.51837 | -6.85588 | 7.09E-12 | 2.62E-09 | 39.44292 | 31.12522 | 40.38657 | 187.2967 | 237.1137 |
| Ptpro | -2.46517 | -6.81729 | 9.28E-12 | 3.34E-09 | 104.4404 | 59.13791 | 71.05044 | 355.2179 | 511.5509 |
| Mir142hg | -2.08032 | -6.79511 | 1.08E-11 | 3.81E-09 | 213.8806 | 184.8838 | 218.3866 | 1022.059 | 715.7321 |
| Sall3 | -2.34453 | -6.78788 | 1.14E-11 | 3.91E-09 | 77.77477 | 54.78038 | 78.52943 | 410.1152 | 302.9786 |
| Cd86 | -2.35044 | -6.74917 | 1.49E-11 | 5.00E-09 | 244.9905 | 155.0036 | 201.1849 | 736.2698 | 1310.712 |
| C1qa | -2.59311 | -6.74313 | 1.55E-11 | 5.10E-09 | 177.7709 | 125.7459 | 160.7984 | 1327.223 | 537.8969 |
| Klf2 | -2.73665 | -6.73871 | 1.60E-11 | 5.15E-09 | 177.2154 | 71.588 | 74.04204 | 794.3963 | 643.2807 |
| Bach1 | -2.44178 | -6.6714 | 2.53E-11 | 7.99E-09 | 64.44196 | 38.59527 | 54.59665 | 245.4233 | 327.1291 |
| Hspb1 | -3.29576 | -6.55806 | 5.45E-11 | 1.68E-08 | 11.11068 | 9.337565 | 8.226893 | 119.4824 | 68.06042 |
| Cd83 | -2.27999 | -6.4732 | 9.59E-11 | 2.90E-08 | 191.6593 | 128.2359 | 263.2606 | 1070.498 | 814.5295 |
| Egr1 | -2.28248 | -6.46356 | 1.02E-10 | 3.04E-08 | 207.2142 | 140.686 | 99.47062 | 768.5623 | 684.9952 |
| Birc3 | -2.43538 | -6.42423 | 1.33E-10 | 3.86E-08 | 49.99807 | 48.55534 | 43.37816 | 180.8382 | 333.7156 |
| Jmjd1c | -2.02364 | -6.37704 | 1.81E-10 | 5.16E-08 | 237.2131 | 199.2014 | 323.8404 | 944.5566 | 1115.313 |
| Tekt3 | -2.39411 | -6.33593 | 2.36E-10 | 6.61E-08 | 42.77613 | 38.59527 | 40.38657 | 264.7988 | 160.2713 |
| Mafb | -2.01523 | -6.30864 | 2.82E-10 | 7.63E-08 | 254.9902 | 154.3811 | 193.7059 | 787.9378 | 838.68 |
| Sqstm1 | -2.31278 | -6.30817 | 2.82E-10 | 7.63E-08 | 165.5492 | 93.37565 | 91.24372 | 469.8564 | 693.7772 |
| Nktr | -2.06538 | -6.29574 | 3.06E-10 | 8.12E-08 | 99.99614 | 80.30306 | 94.98322 | 326.1546 | 443.4905 |
| Smndc1 | -2.601 | -6.24832 | 4.15E-10 | 1.08E-07 | 21.1103 | 24.27767 | 18.69748 | 146.931 | 111.9704 |
| Blnk | -2.09636 | -6.24281 | 4.30E-10 | 1.10E-07 | 168.3268 | 143.7985 | 216.8908 | 584.4949 | 924.3044 |
| Rpl10-ps3 | -2.99356 | -6.22138 | 4.93E-10 | 1.22E-07 | 31.66544 | 21.16515 | 12.71429 | 243.8086 | 105.3839 |
| Irs2 | -2.31474 | -6.22115 | 4.94E-10 | 1.22E-07 | 35.55418 | 30.50271 | 38.14287 | 175.9943 | 169.0533 |
| Smad3 | -2.04072 | -6.13427 | 8.56E-10 | 2.09E-07 | 142.7723 | 99.6007 | 154.8152 | 466.6271 | 623.5213 |
| Gpr137b-ps | -2.08659 | -6.09196 | 1.12E-09 | 2.68E-07 | 55.55341 | 61.00543 | 68.80674 | 230.8916 | 294.1966 |
| Whrn | -1.98813 | -6.06958 | 1.28E-09 | 3.02E-07 | 122.773 | 82.17057 | 112.9328 | 440.7931 | 399.5805 |
| Gm15155 | 2.870331 | 6.067977 | 1.30E-09 | 3.02E-07 | 117.7732 | 206.6714 | 171.269 | 19.37552 | 26.34597 |
| Spag9 | -1.91383 | -6.03219 | 1.62E-09 | 3.71E-07 | 97.774 | 108.3158 | 116.6723 | 356.8325 | 454.468 |
| Vps18 | -2.55071 | -5.9833 | 2.19E-09 | 4.94E-07 | 45.5538 | 24.90017 | 37.39497 | 140.4725 | 283.2192 |
| Slc17a9 | -2.38369 | -5.95379 | 2.62E-09 | 5.83E-07 | 40.55399 | 31.12522 | 31.41177 | 226.0477 | 131.7298 |
| Rhoa | -2.10894 | -5.9481 | 2.71E-09 | 5.95E-07 | 59.44215 | 42.3303 | 52.35296 | 226.0477 | 217.3542 |
| Elf2 | -2.17015 | -5.93273 | 2.98E-09 | 6.40E-07 | 79.44138 | 52.91287 | 69.55464 | 232.5062 | 375.4301 |
| Relb | -2.39803 | -5.93005 | 3.03E-09 | 6.40E-07 | 36.10972 | 29.88021 | 45.62186 | 140.4725 | 252.4822 |
| Tnfaip3 | -2.19709 | -5.92909 | 3.05E-09 | 6.40E-07 | 94.99633 | 67.23047 | 48.61346 | 358.4471 | 287.6102 |
| Aptx | -2.97787 | -5.91793 | 3.26E-09 | 6.76E-07 | 13.33282 | 6.225043 | 17.94959 | 90.4191 | 105.3839 |
| Rassf3 | -2.07007 | -5.91469 | 3.32E-09 | 6.80E-07 | 56.10894 | 58.51541 | 54.59665 | 198.5991 | 276.6327 |
| Npas3 | 2.342379 | 5.906422 | 3.50E-09 | 7.06E-07 | 180.5486 | 263.9418 | 261.0169 | 38.75104 | 54.88743 |
| Id2 | -2.90941 | -5.90052 | 3.62E-09 | 7.22E-07 | 26.1101 | 10.58257 | 14.95799 | 171.1504 | 87.8199 |
| Gria4 | -1.8735 | -5.88734 | 3.92E-09 | 7.66E-07 | 119.4398 | 171.8112 | 125.6471 | 494.0758 | 524.7239 |
| Adamts1 | -2.84614 | -5.88498 | 3.98E-09 | 7.66E-07 | 47.77593 | 25.52268 | 11.21849 | 185.6821 | 223.9407 |
| Dpp10 | 2.311295 | 5.884231 | 4.00E-09 | 7.66E-07 | 218.3249 | 429.528 | 351.5127 | 66.1997 | 68.06042 |
| Macrod2 | 2.073575 | 5.88038 | 4.09E-09 | 7.74E-07 | 331.0983 | 560.8764 | 452.4791 | 103.3361 | 109.7749 |
| Gadd45b | -3.48539 | -5.85969 | 4.64E-09 | 8.67E-07 | 5.555341 | 6.225043 | 8.974793 | 38.75104 | 116.3614 |
| Hsp90b1 | -2.398 | -5.84539 | 5.05E-09 | 9.33E-07 | 86.66332 | 37.97277 | 53.10086 | 387.5104 | 237.1137 |
| Alkbh5 | -2.52451 | -5.83459 | 5.39E-09 | 9.83E-07 | 22.7769 | 15.56261 | 24.68068 | 137.2433 | 103.1884 |
| Lncpint | -2.10097 | -5.81542 | 6.05E-09 | 1.09E-06 | 74.9971 | 70.34299 | 65.81515 | 217.9746 | 388.603 |
| Tiparp | -2.41499 | -5.80457 | 6.45E-09 | 1.15E-06 | 23.88797 | 19.92014 | 22.43698 | 127.5555 | 107.5794 |
| Pltp | -3.78943 | -5.79982 | 6.64E-09 | 1.17E-06 | 6.110875 | 11.20508 | 2.243698 | 37.13641 | 147.0983 |
| Ccnl1 | -2.40686 | -5.79834 | 6.70E-09 | 1.17E-06 | 67.21963 | 30.50271 | 33.65547 | 208.2868 | 259.0687 |
| Kras | -2.325 | -5.78318 | 7.33E-09 | 1.26E-06 | 32.22098 | 23.65517 | 35.89917 | 129.1701 | 177.8353 |
| Man1c1 | -1.84624 | -5.75444 | 8.69E-09 | 1.48E-06 | 214.9917 | 145.666 | 225.1177 | 636.1629 | 768.4241 |
| Gm16576 | -3.23745 | -5.73694 | 9.64E-09 | 1.62E-06 | 9.44408 | 3.735026 | 9.722692 | 56.51194 | 87.8199 |
| Zfp691 | -2.11513 | -5.7331 | 9.86E-09 | 1.64E-06 | 37.77632 | 36.72776 | 43.37816 | 158.2334 | 182.2263 |
| B4galt1 | -1.73978 | -5.727 | 1.02E-08 | 1.68E-06 | 142.2167 | 122.6334 | 145.8404 | 468.2417 | 445.686 |
| Kcnb2 | 2.593856 | 5.720389 | 1.06E-08 | 1.73E-06 | 116.1066 | 177.4137 | 192.2101 | 29.06328 | 24.15047 |
| Tut7 | -2.18831 | -5.68283 | 1.32E-08 | 2.13E-06 | 36.66525 | 32.37023 | 29.16808 | 137.2433 | 162.4668 |
| Pcdh9 | 2.556006 | 5.672991 | 1.40E-08 | 2.24E-06 | 143.3278 | 189.8638 | 174.2606 | 40.36567 | 15.36848 |
| Tc2n | -2.15497 | -5.65469 | 1.56E-08 | 2.46E-06 | 33.88758 | 63.49544 | 53.10086 | 232.5062 | 212.9632 |
| Gm10138 | -2.79402 | -5.64797 | 1.62E-08 | 2.53E-06 | 19.44369 | 8.715061 | 11.21849 | 87.18984 | 96.60188 |
| Mef2c | -1.75151 | -5.63302 | 1.77E-08 | 2.74E-06 | 499.9807 | 301.9146 | 417.3279 | 1386.964 | 1350.231 |
| Nrip1 | -1.88876 | -5.62661 | 1.84E-08 | 2.81E-06 | 185.5484 | 104.5807 | 154.8152 | 558.6608 | 540.0924 |
| Man1a | -1.82623 | -5.6208 | 1.90E-08 | 2.88E-06 | 128.8839 | 115.1633 | 145.8404 | 376.208 | 546.6788 |
| Plcl1 | -1.70721 | -5.60452 | 2.09E-08 | 3.13E-06 | 341.6535 | 334.2848 | 463.6976 | 1086.644 | 1394.141 |
| Litaf | -1.81923 | -5.59999 | 2.14E-08 | 3.18E-06 | 142.2167 | 103.9582 | 124.1513 | 374.5934 | 498.3779 |
| Nlgn1 | 2.289128 | 5.594829 | 2.21E-08 | 3.22E-06 | 165.5492 | 298.8021 | 222.874 | 43.59492 | 50.49644 |
| Nrg3 | 2.158651 | 5.594757 | 2.21E-08 | 3.22E-06 | 179.9931 | 268.9219 | 256.5295 | 56.51194 | 48.30094 |
| Bcor | -1.70755 | -5.59273 | 2.24E-08 | 3.22E-06 | 127.7728 | 118.2758 | 124.1513 | 421.4176 | 384.212 |
| Pmp22 | -2.19241 | -5.56677 | 2.60E-08 | 3.68E-06 | 45.5538 | 38.59527 | 56.09245 | 151.7749 | 276.6327 |
| Pag1 | -1.88631 | -5.56644 | 2.60E-08 | 3.68E-06 | 430.5389 | 231.5716 | 440.5127 | 1307.848 | 1409.509 |
| Clip1 | -1.73372 | -5.56026 | 2.69E-08 | 3.76E-06 | 113.8845 | 100.2232 | 104.7059 | 360.0618 | 346.8886 |
| Susd6 | -1.86075 | -5.55914 | 2.71E-08 | 3.76E-06 | 88.32992 | 66.60797 | 92.73952 | 285.7889 | 313.9561 |
| Srsf11 | -2.06739 | -5.54251 | 2.98E-08 | 4.10E-06 | 61.10875 | 41.08529 | 77.03364 | 266.4134 | 232.7227 |
| Gm47283 | -2.03325 | -5.53262 | 3.15E-08 | 4.30E-06 | 257.7678 | 261.4518 | 427.0505 | 876.7423 | 1705.901 |
| Fscn1 | -2.20068 | -5.52617 | 3.27E-08 | 4.42E-06 | 39.44292 | 32.99273 | 62.82355 | 174.3797 | 239.3092 |
| Fam49b | -1.74574 | -5.50947 | 3.60E-08 | 4.82E-06 | 282.2113 | 173.6787 | 244.5631 | 778.2501 | 788.1836 |
| Il1rapl1 | 2.51351 | 5.485497 | 4.12E-08 | 5.39E-06 | 150.5497 | 322.4573 | 223.6219 | 25.83403 | 57.08293 |
| Hsph1 | -2.29947 | -5.48549 | 4.12E-08 | 5.39E-06 | 29.99884 | 19.29763 | 23.93278 | 124.3263 | 116.3614 |
| Chd2 | -1.86188 | -5.48514 | 4.13E-08 | 5.39E-06 | 131.106 | 82.17057 | 112.1849 | 348.7594 | 441.295 |
| Atp6ap2 | -2.30277 | -5.45495 | 4.90E-08 | 6.34E-06 | 59.99768 | 24.90017 | 37.39497 | 175.9943 | 228.3317 |
| Prrc2c | -1.76002 | -5.44382 | 5.21E-08 | 6.65E-06 | 87.21885 | 107.6933 | 121.1597 | 363.291 | 349.0841 |
| Arih1 | -1.70789 | -5.44333 | 5.23E-08 | 6.65E-06 | 124.9952 | 105.8257 | 120.4118 | 348.7594 | 417.1445 |
| Gm46224 | -2.63697 | -5.44129 | 5.29E-08 | 6.67E-06 | 14.44389 | 12.45009 | 12.71429 | 64.58507 | 100.9929 |
| Pmepa1 | -1.88533 | -5.41034 | 6.29E-08 | 7.87E-06 | 170.549 | 131.9709 | 264.7564 | 692.6749 | 702.5592 |
| P3h2 | -1.82728 | -5.39234 | 6.95E-08 | 8.63E-06 | 117.2177 | 75.32303 | 108.4454 | 384.2812 | 327.1291 |
| Lrrtm4 | 2.528496 | 5.379759 | 7.46E-08 | 9.18E-06 | 127.2173 | 249.6242 | 133.874 | 32.29253 | 26.34597 |
| Spag16 | 2.239508 | 5.374594 | 7.68E-08 | 9.28E-06 | 136.1059 | 219.1215 | 186.9748 | 37.13641 | 39.51895 |
| Adgrl3 | 2.37785 | 5.373978 | 7.70E-08 | 9.28E-06 | 126.6618 | 240.2867 | 211.6555 | 29.06328 | 46.10545 |
| Fnip1 | -1.86904 | -5.3735 | 7.72E-08 | 9.28E-06 | 93.88526 | 58.51541 | 82.26893 | 284.1743 | 287.6102 |
| Plpbp | -3.06767 | -5.36588 | 8.06E-08 | 9.55E-06 | 7.221943 | 7.470052 | 6.731094 | 41.98029 | 79.03791 |
| Rreb1 | -1.80215 | -5.36534 | 8.08E-08 | 9.55E-06 | 379.9853 | 244.6442 | 486.1346 | 1283.628 | 1297.539 |
| Nmt1 | -1.88397 | -5.36342 | 8.17E-08 | 9.58E-06 | 51.66467 | 60.38292 | 75.53784 | 227.6624 | 232.7227 |
| Ntng1 | 3.377905 | 5.347279 | 8.93E-08 | 1.04E-05 | 60.55322 | 128.8584 | 97.22692 | 11.30239 | 6.586492 |
| Stat3 | -1.69495 | -5.33923 | 9.33E-08 | 1.08E-05 | 146.661 | 122.0109 | 118.916 | 366.5203 | 472.0319 |
| Tbc1d15 | -2.72398 | -5.33303 | 9.66E-08 | 1.11E-05 | 24.99903 | 11.20508 | 16.45379 | 67.81432 | 166.8578 |
| Icam1 | -2.07367 | -5.32231 | 1.02E-07 | 1.16E-05 | 66.66409 | 36.10525 | 58.33615 | 185.6821 | 267.8507 |
| Gm5905 | -2.22873 | -5.32087 | 1.03E-07 | 1.16E-05 | 83.33012 | 65.98546 | 48.61346 | 437.5638 | 180.0308 |
| Lamp1 | -2.94443 | -5.32012 | 1.04E-07 | 1.16E-05 | 13.33282 | 4.980035 | 8.226893 | 62.97044 | 74.64691 |
| Skil | -2.13788 | -5.31363 | 1.07E-07 | 1.20E-05 | 86.66332 | 39.21777 | 55.34455 | 217.9746 | 316.1516 |
| Gm45740 | 2.8253 | 5.312245 | 1.08E-07 | 1.20E-05 | 89.99653 | 164.3411 | 114.4286 | 9.68776 | 26.34597 |
| Ski | -1.82116 | -5.31075 | 1.09E-07 | 1.20E-05 | 129.995 | 113.9183 | 192.2101 | 440.7931 | 586.1978 |
| Snx20 | -1.97057 | -5.3015 | 1.15E-07 | 1.25E-05 | 36.66525 | 49.80035 | 55.34455 | 174.3797 | 195.3993 |
| Cdh12 | 2.725745 | 5.291956 | 1.21E-07 | 1.31E-05 | 84.99672 | 168.6987 | 114.4286 | 19.37552 | 17.56398 |
| Sipa1l2 | -1.71997 | -5.28943 | 1.23E-07 | 1.31E-05 | 178.882 | 123.8784 | 201.9328 | 539.2853 | 568.6338 |
| 5430431A17Rik | -1.72127 | -5.2892 | 1.23E-07 | 1.31E-05 | 98.88507 | 102.0907 | 139.1093 | 379.4373 | 366.6481 |
| Skint5 | 2.23992 | 5.280583 | 1.29E-07 | 1.36E-05 | 157.2162 | 290.7095 | 178.748 | 48.4388 | 39.51895 |
| Epb41l2 | -1.81009 | -5.27905 | 1.30E-07 | 1.37E-05 | 679.4182 | 338.0199 | 623.7481 | 1764.787 | 2072.55 |
| Retreg1 | -1.77621 | -5.27079 | 1.36E-07 | 1.42E-05 | 174.9932 | 106.4482 | 180.9916 | 500.5343 | 555.4608 |
| Snx24 | -1.70005 | -5.25286 | 1.50E-07 | 1.55E-05 | 238.3241 | 190.4863 | 268.4959 | 600.6411 | 911.1314 |
| Rnf19b | -2.02029 | -5.24981 | 1.52E-07 | 1.57E-05 | 192.2148 | 111.4283 | 86.00843 | 429.4907 | 627.9123 |
| Arl4c | -1.57913 | -5.23562 | 1.64E-07 | 1.68E-05 | 174.4377 | 146.2885 | 181.7395 | 508.6074 | 491.7914 |
| Cotl1 | -1.81353 | -5.232 | 1.68E-07 | 1.70E-05 | 124.9952 | 133.8384 | 213.8992 | 474.7003 | 632.3032 |
| Nell1 | 2.164089 | 5.231606 | 1.68E-07 | 1.70E-05 | 137.2169 | 232.8166 | 216.1429 | 45.20955 | 41.71445 |
| Morn1 | -2.17282 | -5.22354 | 1.76E-07 | 1.76E-05 | 26.1101 | 32.37023 | 30.66387 | 100.1069 | 169.0533 |
| Plek | -2.14954 | -5.19275 | 2.07E-07 | 2.06E-05 | 322.2098 | 110.1833 | 142.8488 | 810.5426 | 893.5674 |
| Ldlrad4 | -1.75969 | -5.19118 | 2.09E-07 | 2.07E-05 | 1009.961 | 535.3537 | 1065.009 | 2832.055 | 3060.523 |
| Rel | -1.9473 | -5.18808 | 2.12E-07 | 2.09E-05 | 80.55245 | 53.53537 | 56.84035 | 190.526 | 302.9786 |
| Astn2 | 2.166773 | 5.186565 | 2.14E-07 | 2.09E-05 | 129.995 | 210.4065 | 212.4034 | 40.36567 | 41.71445 |
| Rassf1 | -2.56222 | -5.17613 | 2.27E-07 | 2.20E-05 | 13.33282 | 11.82758 | 11.21849 | 62.97044 | 81.2334 |
| Gpa33 | -2.69795 | -5.17131 | 2.32E-07 | 2.24E-05 | 12.22175 | 9.337565 | 9.722692 | 53.28268 | 83.4289 |
| Mertk | -1.82625 | -5.15759 | 2.50E-07 | 2.40E-05 | 924.4088 | 405.8728 | 669.3699 | 2173.288 | 2555.559 |
| Mapkapk2 | -2.01489 | -5.14497 | 2.68E-07 | 2.55E-05 | 54.44234 | 44.82031 | 86.00843 | 200.2137 | 298.5876 |
| Ssh2 | -1.78969 | -5.14393 | 2.69E-07 | 2.55E-05 | 469.4263 | 224.7241 | 352.2606 | 1327.223 | 1084.576 |
| Clasp2 | -1.69986 | -5.14057 | 2.74E-07 | 2.57E-05 | 218.3249 | 153.7586 | 270.7396 | 641.0068 | 750.8601 |
| Snx29 | -1.53287 | -5.13725 | 2.79E-07 | 2.57E-05 | 404.4288 | 303.1596 | 430.0421 | 1033.361 | 1161.418 |
| Ccser1 | 2.238816 | 5.136945 | 2.79E-07 | 2.57E-05 | 161.1049 | 347.3574 | 205.6723 | 40.36567 | 61.47393 |
| Rabgef1 | -2.13348 | -5.1368 | 2.79E-07 | 2.57E-05 | 51.66467 | 49.80035 | 75.53784 | 156.6188 | 362.2571 |
| Fos | -2.04028 | -5.136 | 2.81E-07 | 2.57E-05 | 302.2106 | 220.3665 | 115.1765 | 1073.727 | 676.2132 |
| Sult2b1 | -1.95457 | -5.1355 | 2.81E-07 | 2.57E-05 | 37.77632 | 62.87294 | 55.34455 | 172.7651 | 230.5272 |
| Papola | -1.91893 | -5.12485 | 2.98E-07 | 2.70E-05 | 72.77497 | 41.08529 | 59.83195 | 235.7355 | 201.9858 |
| Slc7a8 | -1.71753 | -5.10697 | 3.27E-07 | 2.95E-05 | 151.1053 | 109.5608 | 176.5043 | 410.1152 | 548.8743 |
| Fgf14 | 2.212965 | 5.102236 | 3.36E-07 | 3.01E-05 | 129.4394 | 237.7967 | 160.0505 | 40.36567 | 35.12796 |
| Robo2 | 2.910747 | 5.071577 | 3.95E-07 | 3.51E-05 | 63.88642 | 131.9709 | 95.73112 | 14.53164 | 10.97749 |
| Erbb4 | 2.345637 | 5.069282 | 3.99E-07 | 3.53E-05 | 104.9959 | 216.6315 | 170.5211 | 25.83403 | 39.51895 |
| Plxdc2 | -1.64404 | -5.05993 | 4.19E-07 | 3.69E-05 | 1002.739 | 551.5389 | 913.933 | 2381.574 | 2761.936 |
| Tmem132a | -3.35682 | -5.05723 | 4.25E-07 | 3.72E-05 | 4.999807 | 5.602539 | 2.991598 | 29.06328 | 65.86492 |
| Foxp2 | 2.984814 | 5.05415 | 4.32E-07 | 3.76E-05 | 58.88662 | 132.5934 | 100.2185 | 9.68776 | 15.36848 |
| Grm8 | 2.183438 | 5.045295 | 4.53E-07 | 3.92E-05 | 155.5495 | 329.3048 | 234.0925 | 38.75104 | 68.06042 |
| Neurl3 | -1.75821 | -5.03534 | 4.77E-07 | 4.10E-05 | 152.7719 | 97.11068 | 111.437 | 327.7692 | 489.5959 |
| Nrxn1 | 1.945983 | 5.034262 | 4.80E-07 | 4.10E-05 | 191.1037 | 319.3447 | 245.311 | 54.89731 | 76.84241 |
| Sgcd | 1.997531 | 5.033316 | 4.82E-07 | 4.10E-05 | 162.216 | 260.8293 | 236.3362 | 45.20955 | 65.86492 |
| Cmklr1 | -2.15247 | -5.0289 | 4.93E-07 | 4.17E-05 | 58.33108 | 23.03266 | 43.37816 | 205.0576 | 164.6623 |
| Klhl7 | -1.97031 | -5.01279 | 5.36E-07 | 4.51E-05 | 31.10991 | 31.12522 | 38.14287 | 119.4824 | 142.7073 |
| Creb5 | -1.88291 | -5.01065 | 5.42E-07 | 4.54E-05 | 118.3288 | 197.9564 | 137.6135 | 376.208 | 742.0781 |
| Cdk13 | -1.72819 | -5.00916 | 5.47E-07 | 4.55E-05 | 76.10817 | 72.2105 | 75.53784 | 201.8283 | 294.1966 |
| Chd7 | -1.52831 | -4.99906 | 5.76E-07 | 4.76E-05 | 228.3245 | 165.5862 | 209.4118 | 602.2558 | 557.6563 |
| Gnai2 | -1.63663 | -4.99833 | 5.78E-07 | 4.76E-05 | 235.5465 | 155.0036 | 242.3194 | 739.499 | 570.8293 |
| Plekhm2 | -1.79433 | -4.99076 | 6.01E-07 | 4.92E-05 | 75.55264 | 112.6733 | 118.916 | 276.1012 | 434.7085 |
| Lpin2 | -1.70886 | -4.98881 | 6.08E-07 | 4.94E-05 | 163.327 | 93.37565 | 154.0673 | 440.7931 | 454.468 |
| Cntnap2 | 1.828463 | 4.982787 | 6.27E-07 | 5.07E-05 | 311.6546 | 608.1867 | 430.79 | 117.8678 | 136.1208 |
| Slc29a3 | -1.53872 | -4.97918 | 6.39E-07 | 5.12E-05 | 203.3255 | 165.5862 | 163.0421 | 573.1925 | 456.6635 |
| Tenm2 | 1.865356 | 4.978954 | 6.39E-07 | 5.12E-05 | 188.8816 | 305.0271 | 278.9665 | 71.04358 | 70.25592 |
| Gm20754 | 3.05432 | 4.975538 | 6.51E-07 | 5.18E-05 | 56.66448 | 103.9582 | 90.49582 | 12.91701 | 6.586492 |
| Myo3a | 3.333502 | 4.974115 | 6.55E-07 | 5.19E-05 | 61.10875 | 100.8457 | 62.07565 | 8.073134 | 6.586492 |
| Baiap2 | -2.1799 | -4.9722 | 6.62E-07 | 5.22E-05 | 20.55476 | 18.67513 | 25.42858 | 85.57522 | 109.7749 |
| Gm15772 | -3.12631 | -4.9596 | 7.06E-07 | 5.54E-05 | 13.88835 | 6.847548 | 4.487396 | 114.6385 | 32.93246 |
| Ildr1 | -2.49976 | -4.95278 | 7.32E-07 | 5.71E-05 | 11.11068 | 12.45009 | 11.21849 | 69.42895 | 61.47393 |
| Erbin | -1.7282 | -4.94661 | 7.55E-07 | 5.86E-05 | 91.10759 | 74.07802 | 91.99162 | 222.8185 | 346.8886 |
| Gm35769 | -2.64342 | -4.94434 | 7.64E-07 | 5.90E-05 | 12.22175 | 10.58257 | 10.47059 | 93.64835 | 43.90995 |
| Naaladl2 | 1.895103 | 4.941829 | 7.74E-07 | 5.95E-05 | 207.2142 | 384.7077 | 276.7228 | 72.6582 | 83.4289 |
| Zc3h12a | -2.35817 | -4.93452 | 8.03E-07 | 6.14E-05 | 14.99942 | 20.54264 | 17.20169 | 62.97044 | 118.5569 |
| Lrrc4c | 2.010336 | 4.932759 | 8.11E-07 | 6.17E-05 | 188.3261 | 341.1324 | 235.5883 | 46.82417 | 81.2334 |
| H2-K2 | -2.63389 | -4.92868 | 8.28E-07 | 6.27E-05 | 12.77728 | 6.225043 | 13.46219 | 59.74119 | 74.64691 |
| Csmd3 | -1.33203 | -4.9262 | 8.38E-07 | 6.31E-05 | 723.3054 | 656.7421 | 703.0254 | 1705.046 | 1791.526 |
| Gpc5 | 1.79461 | 4.907527 | 9.22E-07 | 6.91E-05 | 209.9919 | 314.3647 | 268.4959 | 85.57522 | 65.86492 |
| Lmna | -2.41934 | -4.90352 | 9.41E-07 | 7.02E-05 | 12.77728 | 31.74772 | 16.45379 | 82.34596 | 136.1208 |
| Ncf2 | -1.5786 | -4.89917 | 9.62E-07 | 7.14E-05 | 147.2165 | 112.0508 | 157.8068 | 453.7101 | 375.4301 |
| Ctnna2 | 1.89671 | 4.897506 | 9.71E-07 | 7.17E-05 | 167.2158 | 280.7495 | 231.1009 | 58.12656 | 63.66942 |
| Prpf38b | -1.72153 | -4.88135 | 1.05E-06 | 7.74E-05 | 118.8843 | 69.09798 | 106.2017 | 351.9886 | 294.1966 |
| Arl5c | -2.70948 | -4.87552 | 1.09E-06 | 7.94E-05 | 27.22117 | 8.092557 | 21.68908 | 185.6821 | 61.47393 |
| Nkain2 | 1.831698 | 4.871646 | 1.11E-06 | 8.05E-05 | 183.3263 | 278.8819 | 275.227 | 75.88746 | 61.47393 |
| Ube2h | -1.55111 | -4.86268 | 1.16E-06 | 8.39E-05 | 112.2179 | 150.0235 | 119.6639 | 353.6033 | 392.994 |
| Slc2a1 | -4.32454 | -4.8541 | 1.21E-06 | 8.72E-05 | 1.666602 | 0 | 4.487396 | 16.14627 | 63.66942 |
| Atp6v1a | -1.86826 | -4.8514 | 1.23E-06 | 8.79E-05 | 55.55341 | 33.61523 | 44.87396 | 166.3066 | 160.2713 |
| Aff4 | -1.68844 | -4.84535 | 1.26E-06 | 9.02E-05 | 76.66371 | 92.13064 | 109.1933 | 242.194 | 355.6706 |
| Cacnb2 | -1.54848 | -4.83332 | 1.34E-06 | 9.54E-05 | 171.66 | 145.0435 | 227.3614 | 532.8268 | 526.9194 |
| Gm12381 | 2.33884 | 4.824406 | 1.40E-06 | 9.93E-05 | 88.32992 | 125.7459 | 97.97482 | 19.37552 | 21.95497 |
| Son | -1.70519 | -4.82349 | 1.41E-06 | 9.93E-05 | 213.3251 | 107.6933 | 179.4959 | 494.0758 | 594.9798 |
| Trib1 | -1.97533 | -4.82189 | 1.42E-06 | 9.97E-05 | 41.10952 | 24.27767 | 39.63867 | 124.3263 | 151.4893 |
| Gm33370 | -2.80622 | -4.80962 | 1.51E-06 | 0.000105 | 10.55515 | 5.602539 | 5.983195 | 51.66806 | 52.69194 |
| Gm43464 | -2.37808 | -4.80841 | 1.52E-06 | 0.000106 | 18.33263 | 14.9401 | 9.722692 | 69.42895 | 81.2334 |
| Ythdf3 | -2.01913 | -4.804 | 1.56E-06 | 0.000107 | 35.55418 | 29.2577 | 64.31935 | 156.6188 | 191.0083 |
| Zfp516 | -1.9723 | -4.80061 | 1.58E-06 | 0.000108 | 31.10991 | 26.76769 | 26.17648 | 100.1069 | 120.7524 |
| Syt16 | 2.329267 | 4.800428 | 1.58E-06 | 0.000108 | 86.10779 | 126.9909 | 132.3782 | 16.14627 | 30.73696 |
| Dock2 | -1.6064 | -4.79575 | 1.62E-06 | 0.000111 | 199.4367 | 121.3883 | 181.7395 | 450.4809 | 570.8293 |
| Cfap54 | 3.121666 | 4.79372 | 1.64E-06 | 0.000111 | 51.10914 | 93.99816 | 79.27733 | 4.84388 | 13.17298 |
| Elmo1 | -1.5065 | -4.79069 | 1.66E-06 | 0.000112 | 868.8553 | 537.2213 | 886.2608 | 2045.732 | 2296.49 |
| Adgrb3 | 2.00331 | 4.79051 | 1.66E-06 | 0.000112 | 119.4398 | 188.6188 | 177.2522 | 37.13641 | 43.90995 |
| Arhgap22 | -1.52746 | -4.78161 | 1.74E-06 | 0.000116 | 251.1014 | 170.5662 | 258.7732 | 592.568 | 715.7321 |
| Zswim4 | -2.1701 | -4.77789 | 1.77E-06 | 0.000118 | 14.44389 | 31.74772 | 27.67228 | 106.5654 | 114.1659 |
| Spata21 | -1.91732 | -4.77659 | 1.78E-06 | 0.000118 | 28.33224 | 32.37023 | 29.91598 | 109.7946 | 118.5569 |
| Dcp2 | -1.88283 | -4.77146 | 1.83E-06 | 0.000121 | 37.22079 | 37.35026 | 30.66387 | 113.0239 | 147.0983 |
| Naa50 | -2.02951 | -4.76922 | 1.85E-06 | 0.000121 | 27.77671 | 29.2577 | 25.42858 | 85.57522 | 140.5118 |
| Ptprj | -1.44907 | -4.76882 | 1.85E-06 | 0.000121 | 613.3097 | 423.9255 | 651.4204 | 1464.466 | 1609.3 |
| Gm3379 | -2.6054 | -4.76758 | 1.86E-06 | 0.000122 | 10.55515 | 19.29763 | 7.478994 | 98.49223 | 52.69194 |
| Nlrp3 | -1.84848 | -4.76633 | 1.88E-06 | 0.000122 | 88.32992 | 54.15788 | 74.04204 | 184.0674 | 338.1066 |
| Morf4l1 | -1.59743 | -4.76483 | 1.89E-06 | 0.000122 | 142.2167 | 93.99816 | 102.4622 | 324.54 | 360.0616 |
| Sh3glb1 | -1.80104 | -4.76205 | 1.92E-06 | 0.000123 | 113.8845 | 56.6479 | 73.29414 | 310.0083 | 256.8732 |
| Ehd4 | -1.70164 | -4.74796 | 2.05E-06 | 0.000131 | 123.3286 | 69.09798 | 83.01683 | 289.0182 | 309.5651 |
| Gm26691 | 2.473639 | 4.747568 | 2.06E-06 | 0.000131 | 73.3305 | 130.7259 | 94.98322 | 14.53164 | 21.95497 |
| Kcnq5 | 2.258048 | 4.74571 | 2.08E-06 | 0.000132 | 84.44118 | 160.6061 | 129.3866 | 25.83403 | 26.34597 |
| Gm10073 | -2.21476 | -4.74194 | 2.12E-06 | 0.000134 | 39.99846 | 29.88021 | 25.42858 | 214.7454 | 79.03791 |
| Gm50346 | -4.06403 | -4.73774 | 2.16E-06 | 0.000136 | 1.111068 | 1.245009 | 2.991598 | 29.06328 | 28.54147 |
| Lrp1b | 1.535082 | 4.730143 | 2.24E-06 | 0.000141 | 421.6504 | 643.047 | 531.0086 | 188.9113 | 177.8353 |
| Ubash3b | -1.67848 | -4.72666 | 2.28E-06 | 0.000143 | 386.0962 | 192.3538 | 398.6304 | 972.0053 | 1113.117 |
| Gab2 | -1.60479 | -4.72503 | 2.30E-06 | 0.000143 | 252.2125 | 145.0435 | 249.0505 | 584.4949 | 726.7096 |
| Zfp936 | 2.101666 | 4.724085 | 2.31E-06 | 0.000143 | 98.88507 | 145.0435 | 129.3866 | 27.44865 | 30.73696 |
| A430093F15Rik | -3.0285 | -4.72121 | 2.34E-06 | 0.000145 | 5.555341 | 14.9401 | 2.991598 | 41.98029 | 87.8199 |
| Cntnap5b | 1.897114 | 4.720128 | 2.36E-06 | 0.000145 | 154.4385 | 281.9945 | 200.437 | 54.89731 | 59.27843 |
| Grasp | -3.58168 | -4.70998 | 2.48E-06 | 0.000152 | 2.777671 | 2.490017 | 4.487396 | 58.12656 | 17.56398 |
| Ubac2 | -1.63294 | -4.70637 | 2.52E-06 | 0.000154 | 123.8841 | 76.56803 | 100.9664 | 277.7158 | 346.8886 |
| Numb | -1.43525 | -4.70322 | 2.56E-06 | 0.000155 | 364.9859 | 267.6769 | 406.8573 | 925.1811 | 948.4549 |
| Pxn | -1.6108 | -4.70285 | 2.57E-06 | 0.000155 | 86.66332 | 66.60797 | 97.97482 | 264.7988 | 245.8957 |
| Tanc2 | -1.536 | -4.69876 | 2.62E-06 | 0.000157 | 872.7441 | 565.2339 | 1087.446 | 2396.106 | 2485.303 |
| Chd4 | -1.75048 | -4.69846 | 2.62E-06 | 0.000157 | 159.4383 | 72.2105 | 118.916 | 374.5934 | 412.7535 |
| Sntg1 | 1.915275 | 4.690183 | 2.73E-06 | 0.000162 | 146.1055 | 264.5643 | 234.0925 | 46.82417 | 68.06042 |
| Sec24a | -1.70388 | -4.69018 | 2.73E-06 | 0.000162 | 56.66448 | 45.44282 | 48.61346 | 163.0773 | 164.6623 |
| Grb2 | -1.49227 | -4.68995 | 2.73E-06 | 0.000162 | 160.5494 | 118.2758 | 125.6471 | 385.8958 | 373.2346 |
| Plekho1 | -1.61321 | -4.683 | 2.83E-06 | 0.000167 | 77.21924 | 67.23047 | 65.81515 | 237.3501 | 191.0083 |
| Rin2 | -1.49695 | -4.67596 | 2.93E-06 | 0.000172 | 241.1018 | 199.2014 | 280.4623 | 553.817 | 803.552 |
| Dusp1 | -1.7602 | -4.6749 | 2.94E-06 | 0.000172 | 117.7732 | 102.0907 | 59.08405 | 364.9056 | 265.6552 |
| Slc16a6 | -1.83697 | -4.6742 | 2.95E-06 | 0.000172 | 116.6622 | 62.87294 | 95.73112 | 227.6624 | 430.3175 |
| Med13l | -1.47022 | -4.66982 | 3.01E-06 | 0.000176 | 166.6602 | 122.0109 | 152.5715 | 393.9689 | 421.5355 |
| Bmp2k | -1.63604 | -4.66703 | 3.06E-06 | 0.000177 | 221.1026 | 158.1161 | 322.3446 | 660.3823 | 792.5746 |
| Ppfia2 | 2.55273 | 4.664043 | 3.10E-06 | 0.000179 | 64.44196 | 115.1633 | 83.01683 | 14.53164 | 15.36848 |
| Pik3r1 | -1.55361 | -4.66174 | 3.14E-06 | 0.00018 | 188.3261 | 125.7459 | 201.9328 | 444.0223 | 566.4383 |
| Pard3 | 2.327958 | 4.653491 | 3.26E-06 | 0.000187 | 77.77477 | 98.97819 | 100.9664 | 20.99015 | 15.36848 |
| Bsg | -2.20108 | -4.65051 | 3.31E-06 | 0.000189 | 21.1103 | 26.76769 | 32.15967 | 174.3797 | 68.06042 |
| Kpna4 | -1.86924 | -4.64276 | 3.44E-06 | 0.000196 | 72.21943 | 40.46278 | 47.11766 | 150.1603 | 241.5047 |
| Foxn3 | -1.48058 | -4.63569 | 3.56E-06 | 0.000201 | 750.5266 | 458.1632 | 768.8406 | 1703.431 | 1975.948 |
| Luzp2 | 2.655871 | 4.635281 | 3.56E-06 | 0.000201 | 63.33089 | 95.24317 | 81.52103 | 17.76089 | 6.586492 |
| Abca13 | 2.706403 | 4.632915 | 3.61E-06 | 0.000203 | 64.99749 | 83.41558 | 68.05884 | 12.91701 | 8.78199 |
| Grm5 | 2.338996 | 4.628979 | 3.67E-06 | 0.000206 | 74.9971 | 143.7985 | 113.6807 | 16.14627 | 28.54147 |
| Optn | -2.133 | -4.62756 | 3.70E-06 | 0.000207 | 21.66583 | 32.37023 | 33.65547 | 77.50208 | 180.0308 |
| Rpl10a-ps1 | -2.6358 | -4.62519 | 3.74E-06 | 0.000208 | 11.11068 | 5.602539 | 8.974793 | 58.12656 | 48.30094 |
| Senp2 | -1.59317 | -4.62118 | 3.82E-06 | 0.000212 | 138.8835 | 92.75315 | 133.874 | 301.9352 | 434.7085 |
| Egfem1 | 2.129376 | 4.616506 | 3.90E-06 | 0.000216 | 89.44099 | 148.7785 | 124.8992 | 30.67791 | 24.15047 |
| Thsd7b | 2.001501 | 4.61069 | 4.01E-06 | 0.000221 | 108.3292 | 198.5789 | 179.4959 | 45.20955 | 35.12796 |
| Plch1 | 2.208632 | 4.60729 | 4.08E-06 | 0.000224 | 87.21885 | 145.666 | 125.6471 | 17.76089 | 35.12796 |
| Otulin | -1.82426 | -4.60178 | 4.19E-06 | 0.000229 | 43.88719 | 36.10525 | 30.66387 | 117.8678 | 144.9028 |
| Gm19951 | -2.85479 | -4.59995 | 4.23E-06 | 0.00023 | 9.999614 | 12.45009 | 2.243698 | 74.27283 | 46.10545 |
| Cx3cr1 | -1.71894 | -4.59301 | 4.37E-06 | 0.000237 | 1839.373 | 816.1032 | 1070.992 | 5060.24 | 3117.606 |
| Smarca5 | -1.90212 | -4.59063 | 4.42E-06 | 0.000239 | 25.55457 | 28.0127 | 27.67228 | 103.3361 | 98.79738 |
| Zfp36l1 | -2.12348 | -4.58819 | 4.47E-06 | 0.000241 | 64.44196 | 29.2577 | 29.91598 | 240.5794 | 118.5569 |
| Ddx3x | -1.96249 | -4.58423 | 4.56E-06 | 0.000245 | 181.6597 | 69.09798 | 71.05044 | 466.6271 | 371.0391 |
| Nyap2 | 2.848036 | 4.580963 | 4.63E-06 | 0.000248 | 57.22001 | 74.07802 | 67.31094 | 11.30239 | 6.586492 |
| Tspan13 | -2.02871 | -4.57168 | 4.84E-06 | 0.000258 | 43.33166 | 24.90017 | 21.68908 | 142.0872 | 103.1884 |
| Nfe2l2 | -1.75891 | -4.57104 | 4.85E-06 | 0.000258 | 207.7698 | 84.66059 | 160.0505 | 568.3486 | 452.2725 |
| Tra2b | -1.49991 | -4.5686 | 4.91E-06 | 0.000259 | 90.55206 | 93.99816 | 83.76473 | 232.5062 | 274.4372 |
| Mrnip | -1.72735 | -4.56723 | 4.94E-06 | 0.000259 | 42.77613 | 52.91287 | 48.61346 | 130.7848 | 188.8128 |
| Dnah7c | 2.054319 | 4.566709 | 4.95E-06 | 0.000259 | 101.1072 | 142.5535 | 149.5799 | 22.60477 | 41.71445 |
| Rab3il1 | -1.70209 | -4.56647 | 4.96E-06 | 0.000259 | 92.7742 | 52.91287 | 65.81515 | 256.7256 | 201.9858 |
| Slc23a2 | -1.61314 | -4.56621 | 4.97E-06 | 0.000259 | 84.99672 | 57.2704 | 69.55464 | 206.6722 | 226.1362 |
| Etv6 | -1.56579 | -4.56377 | 5.02E-06 | 0.000261 | 205.5476 | 136.951 | 223.6219 | 448.8662 | 669.6267 |
| Prr16 | 2.236165 | 4.563592 | 5.03E-06 | 0.000261 | 81.66351 | 90.26313 | 104.7059 | 19.37552 | 19.75948 |
| Smo | -2.28981 | -4.56055 | 5.10E-06 | 0.000263 | 17.77709 | 10.58257 | 17.20169 | 90.4191 | 57.08293 |
| Smap2 | -1.55598 | -4.55964 | 5.12E-06 | 0.000263 | 337.2092 | 174.9237 | 233.3446 | 770.1769 | 691.5817 |
| Nfkb1 | -1.6778 | -4.55939 | 5.13E-06 | 0.000263 | 187.215 | 105.2032 | 124.8992 | 337.457 | 555.4608 |
| Calr | -1.95531 | -4.55827 | 5.16E-06 | 0.000264 | 92.21866 | 39.21777 | 80.02523 | 358.4471 | 186.6173 |
| Abl2 | -1.5961 | -4.55641 | 5.20E-06 | 0.000265 | 103.8849 | 108.9383 | 106.2017 | 232.5062 | 412.7535 |
| Gm9794 | -2.24825 | -4.5559 | 5.22E-06 | 0.000265 | 51.10914 | 24.27767 | 24.68068 | 230.8916 | 85.6244 |
| Phldb1 | -2.07977 | -4.55497 | 5.24E-06 | 0.000265 | 17.22156 | 38.59527 | 26.92438 | 95.26298 | 138.3163 |
| Gm22692 | -3.23912 | -4.55138 | 5.33E-06 | 0.000269 | 2.777671 | 4.35753 | 3.739497 | 37.13641 | 30.73696 |
| Pitpnc1 | -1.3507 | -4.5492 | 5.38E-06 | 0.000271 | 248.3237 | 243.3992 | 272.2354 | 566.734 | 733.2961 |
| Bclaf1 | -1.56493 | -4.54732 | 5.43E-06 | 0.000272 | 84.99672 | 65.98546 | 71.05044 | 235.7355 | 201.9858 |
| Dclre1c | -1.82236 | -4.54532 | 5.49E-06 | 0.000274 | 61.66429 | 33.61523 | 57.58825 | 148.5457 | 212.9632 |
| Rtn1 | -1.53236 | -4.54361 | 5.53E-06 | 0.000274 | 118.8843 | 81.54807 | 122.6555 | 300.3206 | 322.7381 |
| Akap6 | 2.189026 | 4.542834 | 5.55E-06 | 0.000274 | 91.66313 | 107.6933 | 115.9244 | 14.53164 | 32.93246 |
| Cebpz | -1.75459 | -4.54282 | 5.55E-06 | 0.000274 | 71.6639 | 39.21777 | 74.78994 | 200.2137 | 217.3542 |
| Dusp6 | -1.67039 | -4.54227 | 5.57E-06 | 0.000274 | 88.88546 | 57.8929 | 84.51263 | 293.8621 | 195.3993 |
| Top1 | -1.5139 | -4.54098 | 5.60E-06 | 0.000275 | 124.4396 | 93.37565 | 88.25213 | 287.4036 | 296.3921 |
| Dcc | 1.786545 | 4.539975 | 5.63E-06 | 0.000275 | 143.8833 | 219.1215 | 193.7059 | 46.82417 | 61.47393 |
| Tpm3-rs7 | -2.92744 | -4.53955 | 5.64E-06 | 0.000275 | 4.444273 | 4.980035 | 7.478994 | 53.28268 | 30.73696 |
| Slc38a10 | -1.5464 | -4.52485 | 6.04E-06 | 0.000294 | 80.55245 | 70.34299 | 84.51263 | 200.2137 | 259.0687 |
| Lpcat3 | -1.6148 | -4.51933 | 6.20E-06 | 0.000301 | 53.33127 | 60.38292 | 69.55464 | 167.9212 | 206.3768 |
| Slc8a1 | -1.46717 | -4.51834 | 6.23E-06 | 0.000301 | 582.7553 | 343.6224 | 605.7985 | 1433.789 | 1389.75 |
| Chn2 | -1.47686 | -4.51776 | 6.25E-06 | 0.000301 | 185.5484 | 138.8185 | 163.79 | 537.6707 | 366.6481 |
| Pfkfb3 | -1.76448 | -4.51553 | 6.32E-06 | 0.000304 | 94.4408 | 48.55534 | 98.72272 | 227.6624 | 320.5426 |
| Malrd1 | 2.186679 | 4.507745 | 6.55E-06 | 0.000314 | 83.33012 | 163.0961 | 133.1261 | 19.37552 | 37.32346 |
| 5033421B08Rik | -1.88059 | -4.50261 | 6.71E-06 | 0.00032 | 36.10972 | 27.39019 | 27.67228 | 93.64835 | 131.7298 |
| Nfatc2 | -1.54644 | -4.5021 | 6.73E-06 | 0.00032 | 89.99653 | 71.588 | 74.04204 | 203.443 | 256.8732 |
| Gabrb1 | 2.455532 | 4.499432 | 6.81E-06 | 0.000323 | 69.44176 | 91.50814 | 71.79834 | 11.30239 | 17.56398 |
| Cnbd1 | 3.367457 | 4.498805 | 6.83E-06 | 0.000323 | 43.88719 | 78.43555 | 51.60506 | 4.84388 | 6.586492 |
| Slc16a1 | -3.08144 | -4.49707 | 6.89E-06 | 0.000324 | 4.444273 | 5.602539 | 3.739497 | 24.2194 | 54.88743 |
| Dgkb | 1.856916 | 4.497024 | 6.89E-06 | 0.000324 | 137.7725 | 265.8094 | 201.9328 | 48.4388 | 63.66942 |
| Trp53inp2 | -2.21474 | -4.4945 | 6.97E-06 | 0.000326 | 17.22156 | 14.3176 | 17.20169 | 53.28268 | 98.79738 |
| Senp6 | -1.55929 | -4.49433 | 6.98E-06 | 0.000326 | 61.66429 | 62.25043 | 60.57985 | 182.4528 | 180.0308 |
| Prkcd | -1.50987 | -4.48952 | 7.14E-06 | 0.000332 | 193.8814 | 122.6334 | 173.5127 | 393.9689 | 537.8969 |
| Rffl | -1.39066 | -4.48376 | 7.33E-06 | 0.000341 | 144.4389 | 146.2885 | 180.9916 | 387.5104 | 436.904 |
| A330008L17Rik | 3.168403 | 4.482156 | 7.39E-06 | 0.000342 | 48.887 | 75.94553 | 50.85716 | 6.458507 | 6.586492 |
| Akap8l | -1.58767 | -4.47825 | 7.53E-06 | 0.000346 | 71.6639 | 52.91287 | 70.30254 | 203.443 | 186.6173 |
| Slc25a21 | 1.798134 | 4.478126 | 7.53E-06 | 0.000346 | 126.6618 | 174.3012 | 159.3026 | 40.36567 | 48.30094 |
| Pde1c | 1.937071 | 4.473586 | 7.69E-06 | 0.000353 | 103.8849 | 190.4863 | 175.0085 | 41.98029 | 39.51895 |
| Mtss1 | -1.3738 | -4.4719 | 7.75E-06 | 0.000355 | 138.328 | 141.3085 | 130.8824 | 372.9788 | 335.9111 |
| E230029C05Rik | -1.54799 | -4.46598 | 7.97E-06 | 0.000363 | 240.5463 | 127.6134 | 228.8572 | 566.734 | 597.1753 |
| Nfia | -1.45716 | -4.45573 | 8.36E-06 | 0.00038 | 243.8795 | 147.5335 | 227.3614 | 568.3486 | 564.2428 |
| Purb | -2.04536 | -4.45019 | 8.58E-06 | 0.000389 | 44.44273 | 23.65517 | 35.15127 | 196.9845 | 85.6244 |
| Skint6 | 2.054831 | 4.448323 | 8.65E-06 | 0.000391 | 87.77439 | 161.2286 | 121.1597 | 30.67791 | 28.54147 |
| G3bp1 | -1.69876 | -4.44096 | 8.96E-06 | 0.000403 | 51.66467 | 36.10525 | 45.62186 | 135.6286 | 153.6848 |
| Jak1 | -1.59445 | -4.43818 | 9.07E-06 | 0.000407 | 195.548 | 95.86567 | 165.2858 | 432.72 | 487.4004 |
| Ubald2 | -2.91036 | -4.43789 | 9.08E-06 | 0.000407 | 6.110875 | 3.112522 | 6.731094 | 33.90716 | 46.10545 |
| Tnf | -2.39746 | -4.42655 | 9.58E-06 | 0.000428 | 24.4435 | 7.470052 | 13.46219 | 64.58507 | 96.60188 |
| Plscr4 | -1.80607 | -4.42439 | 9.67E-06 | 0.000431 | 32.77651 | 48.55534 | 35.15127 | 108.18 | 164.6623 |
| Rnf185 | -1.89755 | -4.42038 | 9.85E-06 | 0.000437 | 33.88758 | 28.0127 | 37.39497 | 85.57522 | 162.4668 |
| Gm29536 | 2.310544 | 4.419751 | 9.88E-06 | 0.000437 | 66.10856 | 85.9056 | 100.2185 | 14.53164 | 19.75948 |
| Kdm6b | -2.16123 | -4.41509 | 1.01E-05 | 0.000446 | 26.1101 | 16.80762 | 44.12606 | 83.96059 | 175.6398 |
| Zdhhc20 | -1.4828 | -4.41419 | 1.01E-05 | 0.000446 | 76.66371 | 71.588 | 78.52943 | 217.9746 | 204.1813 |
| Dip2b | -1.38366 | -4.41178 | 1.03E-05 | 0.00045 | 406.0954 | 293.8221 | 406.1094 | 799.2402 | 1126.29 |
| Gm15964 | -3.13744 | -4.41053 | 1.03E-05 | 0.000451 | 7.221943 | 1.867513 | 3.739497 | 30.67791 | 46.10545 |
| Ralyl | 1.784271 | 4.406334 | 1.05E-05 | 0.000458 | 133.8837 | 219.744 | 184.7311 | 43.59492 | 61.47393 |
| Trim26 | -1.76898 | -4.40622 | 1.05E-05 | 0.000458 | 64.44196 | 32.99273 | 48.61346 | 182.4528 | 149.2938 |
| Rtl4 | 2.522329 | 4.402639 | 1.07E-05 | 0.000464 | 53.33127 | 105.2032 | 79.27733 | 16.14627 | 10.97749 |
| Khdrbs2 | 2.025778 | 4.398957 | 1.09E-05 | 0.00047 | 83.88565 | 135.7059 | 136.8656 | 25.83403 | 32.93246 |
| Unc93b1 | -1.66699 | -4.39836 | 1.09E-05 | 0.00047 | 106.107 | 59.76042 | 124.8992 | 343.9155 | 270.0462 |
| Gm32442 | 3.774825 | 4.397862 | 1.09E-05 | 0.00047 | 58.33108 | 59.13791 | 38.14287 | 1.614627 | 6.586492 |
| Nav2 | -1.32917 | -4.39577 | 1.10E-05 | 0.000473 | 1482.721 | 945.5841 | 1393.337 | 3020.967 | 3381.066 |
| Galnt13 | 1.874812 | 4.391517 | 1.13E-05 | 0.000481 | 110.5513 | 183.6388 | 165.2858 | 33.90716 | 50.49644 |
| Tafa2 | 2.225687 | 4.388494 | 1.14E-05 | 0.000486 | 68.88623 | 126.3684 | 93.48742 | 19.37552 | 21.95497 |
| Gla | -3.47999 | -4.38803 | 1.14E-05 | 0.000486 | 4.999807 | 1.245009 | 2.991598 | 17.76089 | 52.69194 |
| Fam168a | -1.56493 | -4.38605 | 1.15E-05 | 0.000489 | 119.9954 | 67.23047 | 103.958 | 295.4767 | 278.8282 |
| Tead2 | -1.95483 | -4.38305 | 1.17E-05 | 0.000495 | 25.55457 | 19.29763 | 28.42018 | 75.88746 | 114.1659 |
| Agmo | -1.35534 | -4.38229 | 1.17E-05 | 0.000495 | 149.9942 | 129.4809 | 156.311 | 389.125 | 353.4751 |
| Trmt10b | -2.34752 | -4.37889 | 1.19E-05 | 0.000502 | 7.777477 | 14.3176 | 20.19328 | 54.89731 | 87.8199 |
| Fbxo11 | -1.49462 | -4.37814 | 1.20E-05 | 0.000502 | 103.8849 | 85.9056 | 97.97482 | 219.5892 | 322.7381 |
| Pdgfb | -1.49234 | -4.37203 | 1.23E-05 | 0.000514 | 189.4371 | 172.4337 | 204.1765 | 374.5934 | 689.3862 |
| Gm7135 | 2.655606 | 4.371362 | 1.23E-05 | 0.000514 | 74.44157 | 68.47548 | 50.85716 | 9.68776 | 10.97749 |
| Snx9 | -1.56954 | -4.37099 | 1.24E-05 | 0.000514 | 51.10914 | 56.6479 | 52.35296 | 156.6188 | 160.2713 |
| Nav3 | -1.33489 | -4.36535 | 1.27E-05 | 0.000526 | 442.2051 | 370.3901 | 557.185 | 1267.482 | 1034.079 |
| Brd1 | -1.7057 | -4.36496 | 1.27E-05 | 0.000526 | 54.44234 | 39.84028 | 35.15127 | 135.6286 | 147.0983 |
| Zeb2 | -1.42625 | -4.36271 | 1.28E-05 | 0.00053 | 531.6461 | 299.4246 | 487.6304 | 1264.253 | 1097.749 |
| Arvcf | -3.74516 | -4.35685 | 1.32E-05 | 0.000543 | 2.222136 | 1.867513 | 1.495799 | 24.2194 | 26.34597 |
| 9830132P13Rik | -1.76599 | -4.35158 | 1.35E-05 | 0.000554 | 43.88719 | 72.83301 | 47.86556 | 135.6286 | 239.3092 |
| Sgcz | 1.867008 | 4.34116 | 1.42E-05 | 0.00058 | 122.2175 | 246.5117 | 183.2353 | 40.36567 | 61.47393 |
| Map3k2 | -1.79817 | -4.33523 | 1.46E-05 | 0.000592 | 48.33147 | 28.6352 | 32.90757 | 111.4092 | 144.9028 |
| N4bp2l2 | -1.46766 | -4.33479 | 1.46E-05 | 0.000592 | 100.5517 | 78.43555 | 89.74793 | 217.9746 | 278.8282 |
| Rpl13a-ps1 | -2.33123 | -4.3347 | 1.46E-05 | 0.000592 | 26.66564 | 18.67513 | 8.226893 | 121.097 | 59.27843 |
| Naa20 | -2.43356 | -4.3341 | 1.46E-05 | 0.000592 | 6.666409 | 11.20508 | 11.21849 | 45.20955 | 59.27843 |
| Pip5k1c | -2.48284 | -4.33367 | 1.47E-05 | 0.000592 | 23.33243 | 6.847548 | 8.226893 | 66.1997 | 79.03791 |
| Tnrc18 | -1.46614 | -4.33131 | 1.48E-05 | 0.000597 | 97.21847 | 78.43555 | 97.97482 | 219.5892 | 285.4147 |
| Nfkbia | -1.48872 | -4.32802 | 1.50E-05 | 0.000601 | 341.6535 | 183.0163 | 207.1681 | 713.665 | 656.4537 |
| Qk | -1.38057 | -4.32768 | 1.51E-05 | 0.000601 | 549.4232 | 314.9872 | 453.9749 | 1125.395 | 1163.614 |
| Trp63 | 2.747018 | 4.327512 | 1.51E-05 | 0.000601 | 46.10933 | 77.81304 | 67.31094 | 6.458507 | 13.17298 |
| Stk32a | 3.404864 | 4.326855 | 1.51E-05 | 0.000601 | 46.10933 | 54.15788 | 49.36136 | 3.229253 | 6.586492 |
| Lyn | -1.46479 | -4.32666 | 1.51E-05 | 0.000601 | 604.4211 | 333.0398 | 575.1346 | 1181.907 | 1602.713 |
| Pdcd4 | -1.52139 | -4.31731 | 1.58E-05 | 0.000624 | 66.10856 | 72.2105 | 97.22692 | 206.6722 | 243.7002 |
| Gm13166 | -3.68903 | -4.31668 | 1.58E-05 | 0.000624 | 3.333205 | 0.622504 | 2.243698 | 27.44865 | 26.34597 |
| Opcml | 1.603804 | 4.3164 | 1.59E-05 | 0.000624 | 172.2156 | 234.6841 | 236.3362 | 80.73134 | 59.27843 |
| Rcsd1 | -1.64577 | -4.31618 | 1.59E-05 | 0.000624 | 156.6606 | 73.45551 | 163.79 | 382.6665 | 439.0995 |
| Cntn4 | 1.82429 | 4.315767 | 1.59E-05 | 0.000624 | 152.2163 | 308.1397 | 187.7227 | 48.4388 | 74.64691 |
| Lama2 | 2.134868 | 4.314703 | 1.60E-05 | 0.000625 | 86.66332 | 218.499 | 178.748 | 50.05343 | 21.95497 |
| Mbnl2 | -1.42215 | -4.30969 | 1.63E-05 | 0.000638 | 154.4385 | 103.9582 | 151.0757 | 345.5301 | 386.4075 |
| Gm45075 | -2.01789 | -4.30661 | 1.66E-05 | 0.000645 | 16.11049 | 28.6352 | 17.94959 | 92.03372 | 76.84241 |
| Kansl1 | -1.24952 | -4.30514 | 1.67E-05 | 0.000648 | 262.2121 | 227.8366 | 240.8236 | 568.3486 | 590.5888 |
| Tgfbr2 | -1.6177 | -4.30452 | 1.67E-05 | 0.000648 | 368.8746 | 151.2686 | 217.6387 | 781.4793 | 728.9051 |
| Unc13c | 1.907598 | 4.30076 | 1.70E-05 | 0.000656 | 94.99633 | 173.0562 | 137.6135 | 38.75104 | 32.93246 |
| Fli1 | -1.48198 | -4.30067 | 1.70E-05 | 0.000656 | 242.7684 | 139.441 | 264.0085 | 579.651 | 623.5213 |
| Arpc2 | -1.71517 | -4.29638 | 1.74E-05 | 0.000667 | 142.7723 | 57.2704 | 114.4286 | 298.7059 | 390.7985 |
| Btg2 | -1.85023 | -4.29413 | 1.75E-05 | 0.000672 | 194.4369 | 79.05805 | 79.27733 | 527.9829 | 320.5426 |
| Map4k4 | -1.44652 | -4.2893 | 1.79E-05 | 0.000685 | 279.4337 | 160.6061 | 259.5211 | 565.1194 | 706.9502 |
| Arid1b | -1.43702 | -4.28785 | 1.80E-05 | 0.000688 | 172.2156 | 141.3085 | 236.3362 | 439.1785 | 553.2653 |
| Atxn7l1 | -1.43637 | -4.28498 | 1.83E-05 | 0.000695 | 78.88584 | 80.92557 | 97.97482 | 221.2039 | 243.7002 |
| Ptpra | -1.40439 | -4.28084 | 1.86E-05 | 0.000706 | 96.1074 | 89.64063 | 83.76473 | 232.5062 | 243.7002 |
| Map2k2 | -1.49017 | -4.27308 | 1.93E-05 | 0.00073 | 61.66429 | 68.47548 | 73.29414 | 172.7651 | 208.5723 |
| Nfic | -1.87042 | -4.27178 | 1.94E-05 | 0.000732 | 23.33243 | 24.90017 | 37.39497 | 121.097 | 85.6244 |
| Ggnbp2 | -1.57337 | -4.27016 | 1.95E-05 | 0.000735 | 69.9973 | 52.91287 | 93.48742 | 219.5892 | 208.5723 |
| Frmd4b | -1.37811 | -4.26934 | 1.96E-05 | 0.000735 | 317.7655 | 206.0489 | 346.2774 | 734.6552 | 772.8151 |
| Inpp4b | -1.36865 | -4.26924 | 1.96E-05 | 0.000735 | 232.2133 | 212.274 | 317.8572 | 568.3486 | 744.2736 |
| Car10 | 2.031128 | 4.264787 | 2.00E-05 | 0.000748 | 91.10759 | 160.6061 | 133.1261 | 43.59492 | 17.56398 |
| Rheb | -1.65124 | -4.26418 | 2.01E-05 | 0.000748 | 48.887 | 34.23774 | 50.85716 | 135.6286 | 144.9028 |
| Slmap | -1.56751 | -4.26319 | 2.02E-05 | 0.00075 | 112.7734 | 62.25043 | 77.78154 | 230.8916 | 270.0462 |
| Cntnap5c | 1.916891 | 4.254541 | 2.09E-05 | 0.000777 | 103.3293 | 261.4518 | 214.6471 | 51.66806 | 50.49644 |
| Gm12932 | -1.59766 | -4.25386 | 2.10E-05 | 0.000778 | 44.44273 | 57.2704 | 53.10086 | 179.2236 | 131.7298 |
| A230004M16Rik | -1.72089 | -4.25051 | 2.13E-05 | 0.000787 | 34.44311 | 61.00543 | 37.39497 | 153.3895 | 138.3163 |
| Dgkd | -1.45854 | -4.24889 | 2.15E-05 | 0.000791 | 164.4381 | 94.62066 | 131.6303 | 345.5301 | 371.0391 |
| Chd9 | -1.32138 | -4.24766 | 2.16E-05 | 0.000794 | 472.204 | 342.9999 | 565.4119 | 1164.146 | 1135.072 |
| Rlim | -1.69608 | -4.24661 | 2.17E-05 | 0.000795 | 63.88642 | 39.21777 | 59.08405 | 132.3994 | 219.5497 |
| Nkain3 | 2.376839 | 4.242809 | 2.21E-05 | 0.000807 | 60.55322 | 115.1633 | 101.7143 | 27.44865 | 6.586492 |
| Lsamp | 1.463115 | 4.241422 | 2.22E-05 | 0.000807 | 364.4304 | 606.9417 | 473.4203 | 148.5457 | 201.9858 |
| Ube2cbp | 3.008875 | 4.240819 | 2.23E-05 | 0.000807 | 43.88719 | 64.11795 | 49.36136 | 6.458507 | 6.586492 |
| Usp36 | -1.95986 | -4.2408 | 2.23E-05 | 0.000807 | 28.33224 | 16.80762 | 40.38657 | 92.03372 | 129.5343 |
| Ralgds | -2.76341 | -4.24068 | 2.23E-05 | 0.000807 | 6.110875 | 4.980035 | 5.235296 | 30.67791 | 43.90995 |
| Dok6 | 2.678008 | 4.23817 | 2.25E-05 | 0.000814 | 44.99826 | 77.81304 | 81.52103 | 4.84388 | 17.56398 |
| Macf1 | -1.26284 | -4.23764 | 2.26E-05 | 0.000814 | 536.0904 | 367.9001 | 494.3615 | 1104.405 | 1132.877 |
| Ptp4a2 | -1.42231 | -4.23697 | 2.27E-05 | 0.000815 | 277.7671 | 161.8511 | 180.2437 | 581.2656 | 526.9194 |
| Add2 | 2.783415 | 4.225754 | 2.38E-05 | 0.000854 | 41.66506 | 70.34299 | 63.57145 | 6.458507 | 10.97749 |
| Eml4 | -1.38127 | -4.22347 | 2.41E-05 | 0.00086 | 99.99614 | 107.6933 | 126.395 | 259.9549 | 320.5426 |
| Ubxn4 | -1.46392 | -4.22328 | 2.41E-05 | 0.00086 | 100.5517 | 74.07802 | 77.78154 | 206.6722 | 259.0687 |
| Zswim6 | -1.5145 | -4.22178 | 2.42E-05 | 0.000863 | 329.9873 | 177.4137 | 262.5127 | 555.4316 | 913.3269 |
| Usp34 | -1.3011 | -4.22 | 2.44E-05 | 0.000868 | 131.6616 | 129.4809 | 136.8656 | 318.0815 | 335.9111 |
| Zbtb11 | -1.82907 | -4.21669 | 2.48E-05 | 0.000879 | 28.88777 | 26.76769 | 37.39497 | 80.73134 | 140.5118 |
| Gm32296 | -2.43595 | -4.21542 | 2.49E-05 | 0.000882 | 7.221943 | 17.43012 | 6.731094 | 56.51194 | 57.08293 |
| Cmip | -1.36698 | -4.21479 | 2.50E-05 | 0.000882 | 304.9882 | 263.3193 | 272.9833 | 536.0561 | 913.3269 |
| Calm2 | -1.60316 | -4.21015 | 2.55E-05 | 0.000898 | 275.5449 | 120.7658 | 142.1009 | 527.9829 | 564.2428 |
| Zranb2 | -2.11767 | -4.20244 | 2.64E-05 | 0.000927 | 22.7769 | 9.96007 | 20.94118 | 88.80447 | 65.86492 |
| Gm5781 | -1.73181 | -4.20201 | 2.65E-05 | 0.000927 | 91.66313 | 52.29037 | 53.10086 | 284.1743 | 151.4893 |
| Apbb1ip | -1.45053 | -4.20118 | 2.66E-05 | 0.000928 | 206.1032 | 123.8784 | 234.0925 | 532.8268 | 493.9869 |
| Cd300a | -1.5661 | -4.20083 | 2.66E-05 | 0.000928 | 82.21905 | 49.80035 | 57.58825 | 190.526 | 184.4218 |
| Irak2 | -1.39837 | -4.19884 | 2.68E-05 | 0.000934 | 239.9907 | 145.0435 | 214.6471 | 466.6271 | 588.3933 |
| Ophn1 | -1.39793 | -4.19767 | 2.70E-05 | 0.000936 | 331.6539 | 217.254 | 403.1178 | 783.094 | 889.1764 |
| Nipbl | -1.44251 | -4.19152 | 2.77E-05 | 0.00096 | 166.1047 | 98.97819 | 112.9328 | 339.0716 | 346.8886 |
| Pkig | -1.45603 | -4.18802 | 2.81E-05 | 0.000973 | 92.21866 | 77.19054 | 70.30254 | 247.0379 | 191.0083 |
| Abca1 | -1.58742 | -4.18589 | 2.84E-05 | 0.000977 | 84.99672 | 54.15788 | 59.83195 | 161.4627 | 239.3092 |
| Tgfb1 | -1.85984 | -4.18564 | 2.84E-05 | 0.000977 | 77.21924 | 60.38292 | 105.4538 | 148.5457 | 441.295 |
| Slc25a25 | -1.61658 | -4.18543 | 2.85E-05 | 0.000977 | 59.44215 | 45.44282 | 44.12606 | 124.3263 | 182.2263 |
| Thsd7a | 2.385936 | 4.1811 | 2.90E-05 | 0.000993 | 49.99807 | 106.4482 | 75.53784 | 16.14627 | 13.17298 |
| Homer1 | -1.90132 | -4.18071 | 2.91E-05 | 0.000993 | 20.55476 | 23.03266 | 19.44538 | 67.81432 | 90.01539 |
| Veph1 | 2.480706 | 4.179297 | 2.92E-05 | 0.000997 | 46.10933 | 83.41558 | 76.28574 | 9.68776 | 15.36848 |
| Erc2 | 1.794429 | 4.178657 | 2.93E-05 | 0.000997 | 141.6612 | 310.6297 | 204.9244 | 48.4388 | 79.03791 |
| Zdhhc14 | -1.52139 | -4.17755 | 2.95E-05 | 0.001 | 127.2173 | 119.5208 | 139.8572 | 243.8086 | 498.3779 |
| Galnt7 | -1.55436 | -4.17629 | 2.96E-05 | 0.001003 | 63.33089 | 46.06532 | 76.28574 | 174.3797 | 188.8128 |
| Rubcnl | -1.71184 | -4.1751 | 2.98E-05 | 0.001006 | 42.77613 | 31.12522 | 50.85716 | 163.0773 | 107.5794 |
| Ctnna3 | 1.535278 | 4.173568 | 3.00E-05 | 0.001011 | 286.1001 | 586.3991 | 481.6472 | 153.3895 | 158.0758 |
| Asb12 | 3.005823 | 4.172042 | 3.02E-05 | 0.001015 | 38.33185 | 62.25043 | 54.59665 | 8.073134 | 4.390995 |
| Zfand3 | -1.26519 | -4.17131 | 3.03E-05 | 0.001016 | 194.9925 | 164.9637 | 204.1765 | 427.8761 | 476.4229 |
| Nr3c1 | -1.65202 | -4.17083 | 3.03E-05 | 0.001016 | 96.66293 | 48.55534 | 71.05044 | 274.4865 | 177.8353 |
| Hk2 | -1.52992 | -4.16942 | 3.05E-05 | 0.001017 | 159.9938 | 80.92557 | 100.9664 | 314.8522 | 344.6931 |
| Edil3 | 1.921279 | 4.169306 | 3.06E-05 | 0.001017 | 91.10759 | 164.9637 | 108.4454 | 27.44865 | 37.32346 |
| Iqcm | 1.750867 | 4.169251 | 3.06E-05 | 0.001017 | 119.4398 | 242.7767 | 207.1681 | 51.66806 | 61.47393 |
| 6330562C20Rik | -1.7206 | -4.16794 | 3.07E-05 | 0.00102 | 27.77671 | 29.2577 | 35.15127 | 96.8776 | 105.3839 |
| Calml4 | -2.69515 | -4.16751 | 3.08E-05 | 0.00102 | 6.110875 | 4.35753 | 6.731094 | 38.75104 | 35.12796 |
| Csf3r | -1.42519 | -4.1659 | 3.10E-05 | 0.001025 | 264.9898 | 222.8566 | 416.58 | 894.5032 | 722.3186 |
| Polr2g | -1.4021 | -4.16227 | 3.15E-05 | 0.001039 | 96.66293 | 95.24317 | 76.28574 | 255.111 | 217.3542 |
| Dcakd | -1.76562 | -4.16191 | 3.16E-05 | 0.001039 | 38.33185 | 31.12522 | 70.30254 | 155.0042 | 160.2713 |
| Ncam1 | 2.774309 | 4.157696 | 3.21E-05 | 0.001056 | 56.66448 | 52.91287 | 56.84035 | 12.91701 | 2.195497 |
| Tafa4 | 2.364099 | 4.156529 | 3.23E-05 | 0.001059 | 55.55341 | 96.48817 | 62.82355 | 14.53164 | 13.17298 |
| Nuak1 | -1.45614 | -4.15541 | 3.25E-05 | 0.001061 | 195.548 | 102.7132 | 169.0253 | 418.1883 | 436.904 |
| Cry1 | -1.91758 | -4.15504 | 3.25E-05 | 0.001061 | 19.44369 | 32.99273 | 17.94959 | 85.57522 | 92.21089 |
| Wdfy3 | -1.34288 | -4.15428 | 3.26E-05 | 0.001062 | 155.5495 | 112.0508 | 148.832 | 329.3839 | 375.4301 |
| Gm32828 | 2.827542 | 4.153315 | 3.28E-05 | 0.001064 | 46.10933 | 57.8929 | 59.83195 | 3.229253 | 13.17298 |
| Il6ra | -1.42488 | -4.15095 | 3.31E-05 | 0.001071 | 364.9859 | 214.764 | 275.9749 | 933.2542 | 597.1753 |
| Bin1 | -1.47027 | -4.1504 | 3.32E-05 | 0.001071 | 440.5385 | 211.6515 | 425.5547 | 1015.6 | 974.8008 |
| Itga9 | -1.38652 | -4.15007 | 3.32E-05 | 0.001071 | 161.6604 | 138.8185 | 181.7395 | 326.1546 | 515.9419 |
| Srgn | -1.35353 | -4.15 | 3.32E-05 | 0.001071 | 119.4398 | 96.48817 | 111.437 | 253.4964 | 305.1741 |
| Tbc1d9 | -1.50997 | -4.14844 | 3.35E-05 | 0.001076 | 437.7609 | 234.6841 | 254.2858 | 1085.029 | 674.0177 |
| Ptprd | 1.383451 | 4.146101 | 3.38E-05 | 0.001085 | 406.651 | 681.0198 | 585.6052 | 201.8283 | 226.1362 |
| Tmem232 | 2.684205 | 4.145593 | 3.39E-05 | 0.001085 | 49.44254 | 56.6479 | 57.58825 | 6.458507 | 10.97749 |
| C130073E24Rik | 2.783869 | 4.145094 | 3.40E-05 | 0.001085 | 41.66506 | 81.54807 | 54.59665 | 4.84388 | 13.17298 |
| Klf7 | -1.39993 | -4.14386 | 3.42E-05 | 0.001088 | 91.10759 | 82.79308 | 106.9496 | 217.9746 | 276.6327 |
| Mageb18 | 2.070749 | 4.138226 | 3.50E-05 | 0.001113 | 68.3307 | 117.0308 | 86.75633 | 19.37552 | 24.15047 |
| Fgf13 | -1.31809 | -4.13338 | 3.57E-05 | 0.001134 | 135.5503 | 125.1234 | 172.0169 | 351.9886 | 366.6481 |
| Grik2 | 1.792385 | 4.132464 | 3.59E-05 | 0.001137 | 93.88526 | 153.7586 | 141.353 | 35.52179 | 39.51895 |
| Gm9750 | 3.335788 | 4.130316 | 3.62E-05 | 0.001143 | 39.44292 | 62.25043 | 38.89077 | 4.84388 | 4.390995 |
| Gm9843 | -2.08627 | -4.13026 | 3.62E-05 | 0.001143 | 33.33205 | 37.35026 | 18.69748 | 190.526 | 61.47393 |
| Lrch1 | -1.37022 | -4.12943 | 3.64E-05 | 0.001144 | 322.2098 | 196.7114 | 350.0169 | 713.665 | 783.7926 |
| Zmat4 | 1.856133 | 4.125981 | 3.69E-05 | 0.001158 | 90.55206 | 141.931 | 140.6051 | 25.83403 | 43.90995 |
| Gbp7 | -1.97332 | -4.12579 | 3.69E-05 | 0.001158 | 21.1103 | 23.03266 | 17.94959 | 106.5654 | 54.88743 |
| Csf2rb | -1.98815 | -4.12377 | 3.73E-05 | 0.001165 | 37.77632 | 14.3176 | 21.68908 | 95.26298 | 100.9929 |
| Ythdc1 | -1.92953 | -4.12351 | 3.73E-05 | 0.001165 | 41.10952 | 16.80762 | 23.93278 | 109.7946 | 98.79738 |
| Camk1d | -1.27569 | -4.12177 | 3.76E-05 | 0.001166 | 403.3178 | 258.9618 | 351.5127 | 796.011 | 840.8755 |
| 9330158H04Rik | 2.527222 | 4.121762 | 3.76E-05 | 0.001166 | 51.10914 | 89.64063 | 58.33615 | 6.458507 | 17.56398 |
| Btg3 | -3.18274 | -4.12135 | 3.77E-05 | 0.001166 | 2.777671 | 3.735026 | 2.991598 | 37.13641 | 19.75948 |
| Agbl4 | 1.518301 | 4.121337 | 3.77E-05 | 0.001166 | 208.3253 | 332.4173 | 293.9245 | 79.11671 | 116.3614 |
| Ptn | 2.531246 | 4.119232 | 3.80E-05 | 0.001174 | 76.66371 | 97.11068 | 41.13447 | 8.073134 | 17.56398 |
| Dock4 | -1.32953 | -4.11771 | 3.83E-05 | 0.00118 | 796.0804 | 485.5534 | 874.2944 | 1756.714 | 1855.195 |
| Hpgds | -1.41354 | -4.11628 | 3.85E-05 | 0.001185 | 542.2013 | 267.6769 | 481.6472 | 1130.239 | 1163.614 |
| Gabrb3 | 2.449408 | 4.111381 | 3.93E-05 | 0.001208 | 49.44254 | 68.47548 | 65.06725 | 11.30239 | 10.97749 |
| Dnmt3a | -1.23798 | -4.10976 | 3.96E-05 | 0.001212 | 378.3187 | 268.2994 | 353.7564 | 745.9575 | 827.7025 |
| Dach2 | 2.100432 | 4.109616 | 3.96E-05 | 0.001212 | 66.66409 | 136.3285 | 86.00843 | 20.99015 | 24.15047 |
| Edem1 | -1.55546 | -4.09643 | 4.20E-05 | 0.001278 | 162.7715 | 75.94553 | 97.97482 | 303.5498 | 357.8661 |
| Mgat4a | -1.26123 | -4.09637 | 4.20E-05 | 0.001278 | 204.4366 | 183.6388 | 257.2774 | 516.6806 | 513.7464 |
| Kcnip4 | 1.391162 | 4.093485 | 4.25E-05 | 0.001292 | 242.7684 | 333.6623 | 331.3194 | 108.18 | 122.9479 |
| Il1rapl2 | 1.597152 | 4.086937 | 4.37E-05 | 0.001326 | 207.7698 | 456.2957 | 311.1261 | 111.4092 | 103.1884 |

**Supp Table 1. Top 500 differentially expressed genes in microglia from wildtype and mcKO mice.**

Bulk transcriptome data in purified microglia from 3 mcKO and 2 wildtype control mice. Expression levels were adjusted by the levels of the house keeping genes from each sample. P-values were calculated by t-tests between genotypes, and adjusted by Benjamini-Hochberg. p value < 0.05 |log2Fold Change |> 1.0 was used to calculate the difference of gene expression. DEGs were ranked by the statistical significance.
